# A stress-induced block in dicarboxylate uptake and utilization in Salmonella

**DOI:** 10.1101/648782

**Authors:** Steven J. Hersch, Bojana Radan, Bushra Ilyas, Patrick Lavoie, William Wiley Navarre

**Affiliations:** Department of Molecular Genetics, University of Toronto, Toronto, ON, Canada

## Abstract

Bacteria have evolved to sense and respond to their environment by altering gene expression and metabolism to promote growth and survival. In this work we demonstrate that *Salmonella* displays an extensive (>30 hour) lag in growth when subcultured into media where dicarboxylates such as succinate are the sole carbon source. This growth lag is regulated in part by RpoS, the RssB anti-adaptor IraP, translation elongation factor P, and to a lesser degree the stringent response. We also show that small amounts of proline or citrate can trigger early growth in succinate media and that, at least for proline, this effect requires the multifunctional enzyme/regulator PutA. We demonstrate that activation of RpoS results in the repression of *dctA*, encoding the primary dicarboxylate importer, and that constitutive expression of *dctA* induced growth. This dicarboxylate growth lag phenotype is far more severe across multiple *Salmonella* isolates than in its close relative *E. coli*. Replacing 200 nt of the *Salmonella dctA* promoter region with that of *E. coli* was sufficient to eliminate the observed lag in growth. We hypothesize that this *cis*-regulatory divergence might be an adaptation to *Salmonella*’s virulent lifestyle where levels of phagocyte-produced succinate increase in response to bacterial LPS. We found that impairing *dctA* repression had no effect on *Salmonella*’s survival in acidified succinate or in macrophage but propose alternate hypotheses of fitness advantages acquired by repressing dicarboxylate uptake.

**Importance:** Bacteria have evolved to sense and respond to their environment to maximize their chance of survival. By studying differences in the responses of pathogenic bacteria and closely related non-pathogens, we can gain insight into what environments they encounter inside of an infected host. Here we demonstrate that *Salmonella* diverges from its close relative *E. coli* in its response to dicarboxylates such as the metabolite succinate. We show that this is regulated by stress response proteins and ultimately can be attributed to *Salmonella* repressing its import of dicarboxylates. Understanding this phenomenon may reveal a novel aspect of the *Salmonella* virulence cycle, and our characterization of its regulation yields a number of mutant strains that can be used to further study it.

## Introduction

Bacteria must adapt to changing environmental conditions by sensing their surroundings and integrating signals to initiate rapid growth in nutrient rich situations or instigate defence mechanisms and metabolic hibernation in response to stress (1). Pathogens like *Salmonella* have adapted to survive and replicate within their particular host niche, which is reflected in a variety of subtle differences in regulation compared to their non-pathogenic relatives. Understanding the differences in related pathogenic and commensal bacteria *E. coli* and *Salmonella*, has yielded a wealth of insight regarding the challenges pathogens face in the host environment. These bacteria are closely related, yet *Salmonella* has acquired a number of adaptations that accommodate its virulent lifestyle; for example allowing *Salmonella* to invade tissues and survive within host cells such as macrophage, which is important for *Salmonella* virulence (2–6).

Metabolic modulation is an important component in the adaptation to multiple types of stress. The bacterial stringent response, wherein the second messenger molecule, guanosine 5’-disphosphate 3’-diphosphate (ppGpp) is produced by RelA or SpoT in response to amino acid starvation or other cellular stress cues (7–10). Bacteria can also alter gene expression using the general stress response sigma factor RpoS (σ^S^), which has been linked to virulence in a number of pathogenic bacteria by contributing to virulence gene expression and survival within an infected host (11–13). RpoS can be activated in response to a variety of conditions including starvation, hyper-osmolarity and oxidative stress, and can be regulated at all levels of synthesis from transcription to protein degradation where it is recognized by the adaptor RssB (also known as MviA, SprE, or ExpM) and chaperoned to the ClpXP protease (14–18). In response to specific stresses, the anti-adaptors IraP, RssC (IraM in *E. coli*) and IraD can be induced, which impair RssB and thereby rapidly stabilize RpoS (19–21). Strains with reduced *rpoS* activity have been demonstrated to grow faster than wild-type *E*. coli when grown using the dicarboxylate succinate as a sole carbon source (22–24). Furthermore, aerobic growth using succinate relies on the dicarboxylate importer, DctA, which is known to be regulated by the DcuSR two-component system in response to dicarboxylates, as well as by DctR (YhiF) in *E. coli* strains lacking ATP Synthase activity (25–27).

Our previous studies of *Salmonella* translation Elongation Factor P (EF-P), which facilitates the ribosomal synthesis of difficult-to-translate polyproline/glycine motifs, revealed that strains lacking functional EF-P show rapid growth using succinate as a carbon source (28– 31). Here we demonstrate that activation of RpoS, via the antiadaptor IraP or via the stringent response, triggers wild-type *Salmonella* to shut down import of dicarboxylates like succinate, fumarate, and malate. For reasons that remain unclear, Salmonella resumes import and growth on succinate media after a stereotypical lag of approximately 30 hours. The severity of this lag differs between the vast majority of *Salmonella* and *E. coli* strains, and this difference appears to be due almost entirely to differences within the promoter of the dicarboxylate importer DctA. Furthermore, we find that trace amounts of proline or citrate can “license” early *Salmonella* growth on dicarboxylates. In the case of proline, this phenotype depends on the unsual polyfunctional enzyme/regulator PutA. This work adds to a larger body of evidence that proline is an essential signaling metabolite that modulates key aspects of *Salmonella* growth within the animal host.

## Results

### *Salmonella* delays its growth using dicarboxylic acids as a sole carbon source

Our previous work investigating the role of EF-P in *Salmonella* demonstrated that strains deficient in EF-P activity display a ‘hyper-active’ metabolism relative to wild-type when grown under specific nutrient limited conditions (28, 29). Biolog phenotype microarrays (31) revealed that wild-type *Salmonella enterica* Sv. Typhimurim strain 14028s failed to grow under several varied nutrient conditions whereas isogenic strains lacking functional EF-P grew rapidly. Through a series of further growth assays we determined that the underlying cause of this growth defect was due entirely to the fact that many of the varied growth conditions in the Biolog assay employ a base media where succinate provides the sole carbon source.

Earlier literature reported that many wild-type strains of *E. coli* display an extended lag when subcultured on succinate media and that this growth delay was dependent on the sigma factor RpoS (22–24). Accordingly, we tested the growth of strain 14028s in minimal media containing succinate as the sole carbon source. We found that the *efp* (EF-P) mutant, as well as an *rpoS* mutant, were able to grow readily in succinate with minimal lag (Figure 1). In contrast, wild-type *Salmonella* exhibited an extended lag phase where it failed to grow for over 30 hours before initiating exponential (log) growth. The dicarboxylic acid transporter, DctA, was also required for growth and an isogenic *dctA* deletion strain showed no sign of growth by 48 hours.

**Figure 1:**
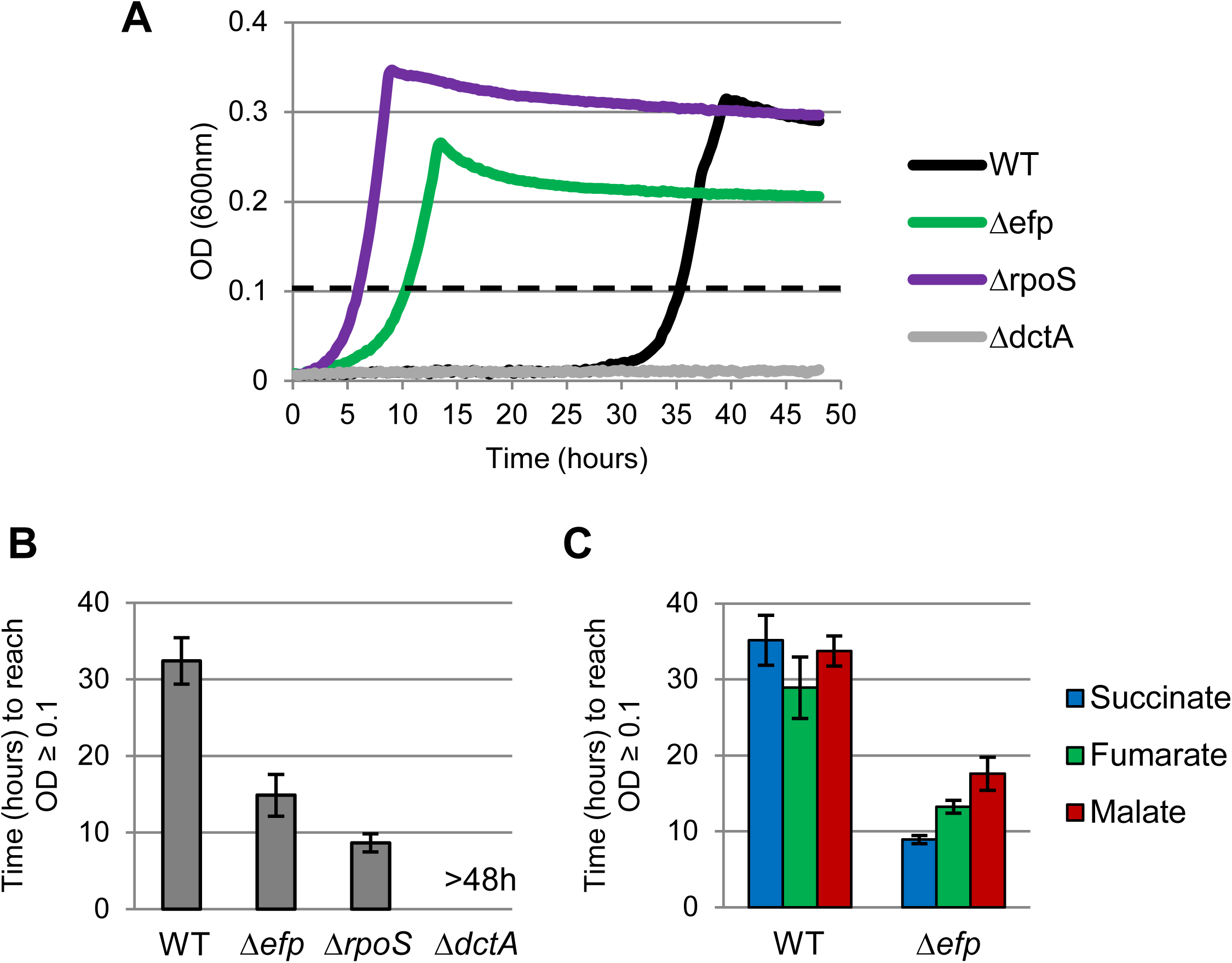
***Salmonella* displays an extended lag phase using dicarboxylic acids as a sole carbon source** but *efp* and *rpoS* mutants do not. **A)** Representative growth curve of *Salmonella* Typhimurium 14028s strains grown in MOPS minimal media with 0.2% succinate as the sole carbon source. Data shows optical density at 600nm (OD_600_) for 48 hours from the time of inoculation. The data shown is representative of greater than three biological replicates. An OD_600_ of 0.1 is emphasized by a dashed line. **B)** Graphs showing the average time in hours that wild-type (WT) or mutant *Salmonella* takes to reach an OD of 0.1 as an analog of the length of lag phase using succinate as the sole carbon source. The data shows the average of at least three biological replicates and error bars show one standard deviation. >48h indicates the strain did not grow by 48 hours. **C)** As in (B) but comparing the use of three different dicarboxylic acids as the sole carbon source.

We note that the wild-type log growth rate, when it finally resumed, was similar to the log growth rate of the *rpoS* mutant, suggesting that the growth delay was due to an extended lag phase upon subculturing into succinate media and not due to slow growth on succinate *per se*. The length of this extended lag phase was remarkably consistent, stereotypically between 30-32 hours across dozens of experiments. Furthermore, colonies of *Salmonella* that grew on succinate media after 30 hours, when recultured, continued to display an extended lag in growth. Occasional colonies that did show rapid growth on succinate were found to have acquired spontaneous mutations in RpoS. Together these results indicate that the bacteria that grew on succinate after 30 hours did not arise from mutation, but rather from a genetically programmed physiological response.

DctA can also import other dicarboxylic acids including malate and fumarate. We therefore tested whether the extended lag of wild-type *Salmonella* would also occur in media with fumarate and malate as the sole carbon source (Figure 1C). We found that, similar to succinate, wild-type strain 14028s displayed a similarly extended lag in growth using either malate or fumarate, while the *efp* mutant grew significantly earlier. This suggests that the growth repression instigated by wild-type *Salmonella* is not specific to succinate but also occurs when subcultured into media using other dicarboxylic acids as the sole carbon source.

These data also indicate that the failure of strain 14028s to grow rapidly on succinate was not due to an inherent inability to utilize this carbon source (i.e. the strain does not lack the factors necessary to import or catabolize succinate), but rather that delayed growth was the result of a regulatory block that prevented the expression of the necessary succinate utilization systems. This regulation involves RpoS, perhaps through a protein(s) that requires *efp* for its efficient translation. We believe this phenotype may have been overlooked in prior studies on *Salmonella* metabolism as many of these studies employed strain LT2, most isolates of which harbor a hypomorphic lesion in the gene encoding RpoS (32).

### Many *Salmonella* but few *E. coli* strains delay growth using dicarboxylates

Prior literature suggested the severity of the growth lag phenotype was much less severe in *E. coli* than what we observed in strain 14028s. To determine if this is indeed the case, or if it was merely a difference in growth conditions, the growth of *Salmonella* 14028s was compared to *E. coli* K12 (strain BW25113). This revealed that, while *E. coli* did display a lag in growth – this lag was much shorter than what was observed for *Salmonella* (Figure 2A).

**Figure 2:**
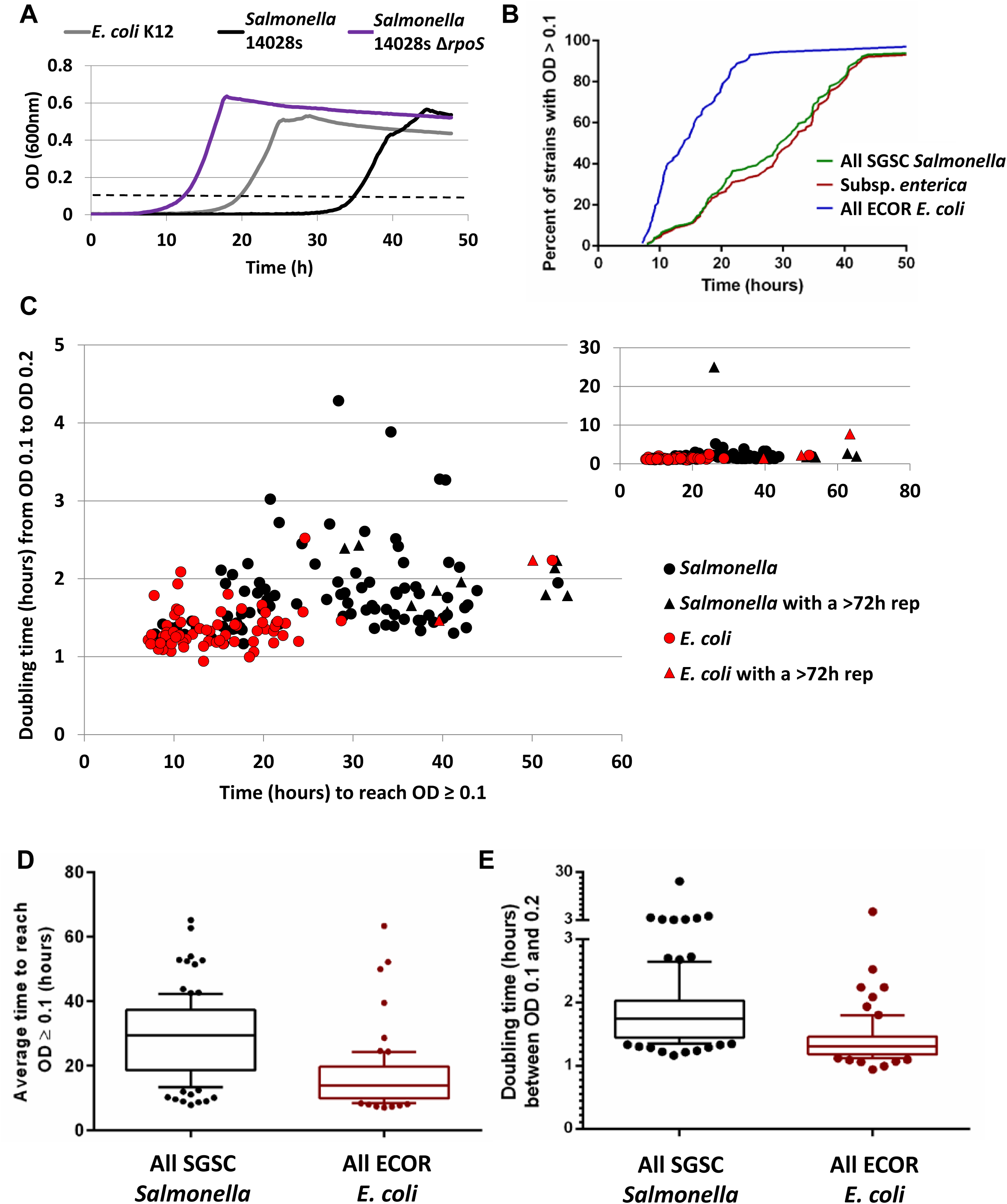
***Salmonella* grows significantly later than *E. coli* using succinate** as the sole carbon source. Growth in MOPS minimal media with 0.2% succinate as the sole carbon source. Growth was conducted in a TECAN Infinite M200 plate reader and reads were taken every 15 minutes. **A)** Representative growth curve. An OD_600_ of 0.1 is emphasized by a dashed line. **B)** Percentage of SGSC *Salmonella* (green line) or ECOR *E. coli* (blue line) strains that (on average) had surpassed an OD_600_ of 0.1 by indicated times post inoculation. Red line is for *Salmonella* but excludes the 14 tested strains of the SARC collection that do not belong to subspecies *enterica*. **C)** Overview of 105 *Salmonella* strains (SGSC collection) and 72 *E. coli strains* (ECOR collection). Each strain is plotted by average time it takes to reach OD 0.1 (x-axis) compared to doubling time during growth from OD 0.1 to OD 0.2. All *Salmonella* are coloured black and all *E. coli* strains are coloured red. Strains with at least one replicate that did not grow by 72 hours are shown as triangles; these values are the average of the remaining replicates. Inset at top right shows zoomed out view to include outliers. **D)** Average time in hours for strains to reach an OD of 0.1. Data compares all 105 *Salmonella* strains to all 72 *E. coli* strains tested. Line indicates the median, boxes show the 25th to 75th percentiles, and whiskers show the 10th to 90th percentiles. An unpaired t-test with Welch’s correction indicated a p-value < 0.0001. **E)** As in (D) but comparing doubling time in hours. An unpaired t-test with Welch’s correction indicated a p-value < 0.05 with all data points included or p-value < 0.0001 when excluding slow growing outliers (doubling time > 5h).

To determine whether the length of the dicarboxylate lag was a characteristic broadly shared among each these two species, or limited to specific strains of *E. coli* and *Salmonella*, growth on succinate media was assessed for all 105 non-typhoidal strains in the *Salmonella* Genetic Stock Centre (SGSC) collection, as well as all 72 strains of the *E. coli* Reference (ECOR) collection. Though there are some exceptions, the lag displayed by the vast majority of *E. coli* strains on succinate media was considerably shorter compared to most *Salmonella* strains (Figure 2B-D). Once logarithmic growth was initiated, *Salmonella* also appeared to trend towards a slightly longer doubling time than the majority of *E. coli* strains (Figure 2C and E).

Hypomorphic variants of *rpoS* accumulate readily in *E. coli* and *Salmonella* during laboratory passage, and there remained the possibility that many strains in the ECOR or SGSC collections acquired such mutations during their cultivation prior to being stored/archived. To ensure that the observed effects were not due to variations in RpoS activity, each strain was also screened for catalase activity as a surrogate indicator of having a functional RpoS. Regardless of catalase activity, the trend was maintained that *E. coli* strains in general showed a shorter lag phase than *Salmonella* when using succinate as the sole carbon source (Figure S1).

### IraP contributes to growth repression in dicarboxylate media

RpoS is activated under specific conditions of stress or nutrient limitation and its expression, synthesis, stability, and activity are regulated at multiple levels. Under ideal growth conditions RpoS protein is destablized through its interaction with RssB, an ‘adaptor’ protein that promotes RpoS degradation via the ClpXP proteases. In response to specific stressors or nutrient limitations, one or more anti-adaptor proteins, IraP, RssC and IraD, can antagonize binding by RssB to alleviate RpoS degradation (14, 19, 20, 33). Systematic deletion of these anti-adaptors revealed that IraP is a primary upstream factor invoved in the dicarboxylate lag, and the rapid growth phenotype of an *iraP* mutant strain could be partially complemented by expressing IraP from its native promoter on a plasmid (Figure 3). In contrast, deletion of the genes encoding the anti-adaptors, RssC and IraD, had comparatively minor effects. These findings demonstrate that, under the laboratory growth conditions we employ, IraP-mediated stabilization of RpoS plays a role in repressing *Salmonella*’s growth using succinate as the sole carbon source.

**Figure 3:**
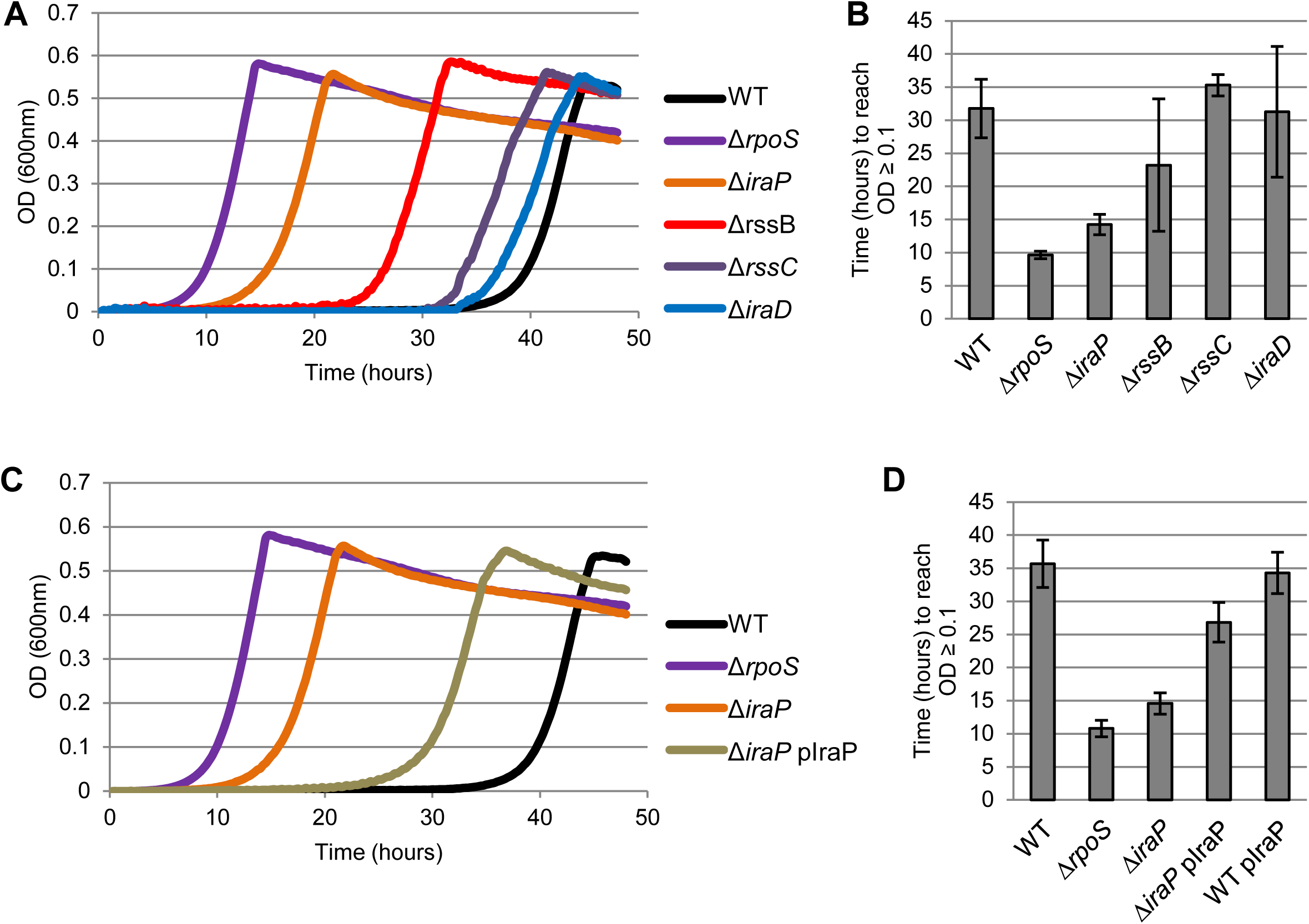
Deletion of the IraP results in early growth on succinate. Growth of *Salmonella* in MOPS minimal media with 0.2% succinate as the sole carbon source. **A)** The three known RssB anti-adaptors were deleted from the *Salmonella* chromosome and growth is shown along with an *rssB* mutant. Data is representative of at least three biological replicates. **B)** Average time in hours for wild-type (WT) or mutant *Salmonella* to reach an OD of 0.1 as an analog of the length of lag phase using succinate as the sole carbon source. The data shows the average of at least three biological replicates and error bars show one standard deviation. **C)** Plasmid expression of the *Salmonella iraP* gene partially complements the growth delay phenotype. **D)** As for B.

### Proline and citrate override the lag observed in succinate media

During these studies it was observed that supplementation of our minimal media with small amounts of LB media (a.k.a. ‘lysogeny broth’ or ‘Luria broth’) would shorten or even eliminate the observed lag phase on succinate (Figure S2). Given that the primary nutrient components in LB are peptides and amino acids, the effects of each amino and and a few other carbon sources in the TCA cycle were examined.

Unlike all other amino acids we found that the addition of 0.005% proline to succinate media could ‘override’ the extended growth lag to enable rapid growth on succinate. The cells did not appear to simply consume the proline as a carbon source since no other individual amino acid showed similar growth induction (Figure S2D). Furthermore, although proline can be used by *Salmonella* as a carbon/nitrogen source, there was no discernible growth on 0.01% proline in the absence of succinate, suggesting the levels we were providing were too small (Figure S2E). A similar early-growth phenomenon could be invoked through the addition of small amounts of citrate. Growth on succinate in the presence of either proline or citrate occurred in a manner resembling a diauxy wherein growth would cease again following exhaustion of the proline or citrate (Figure 4A and B).

**Figure 4:**
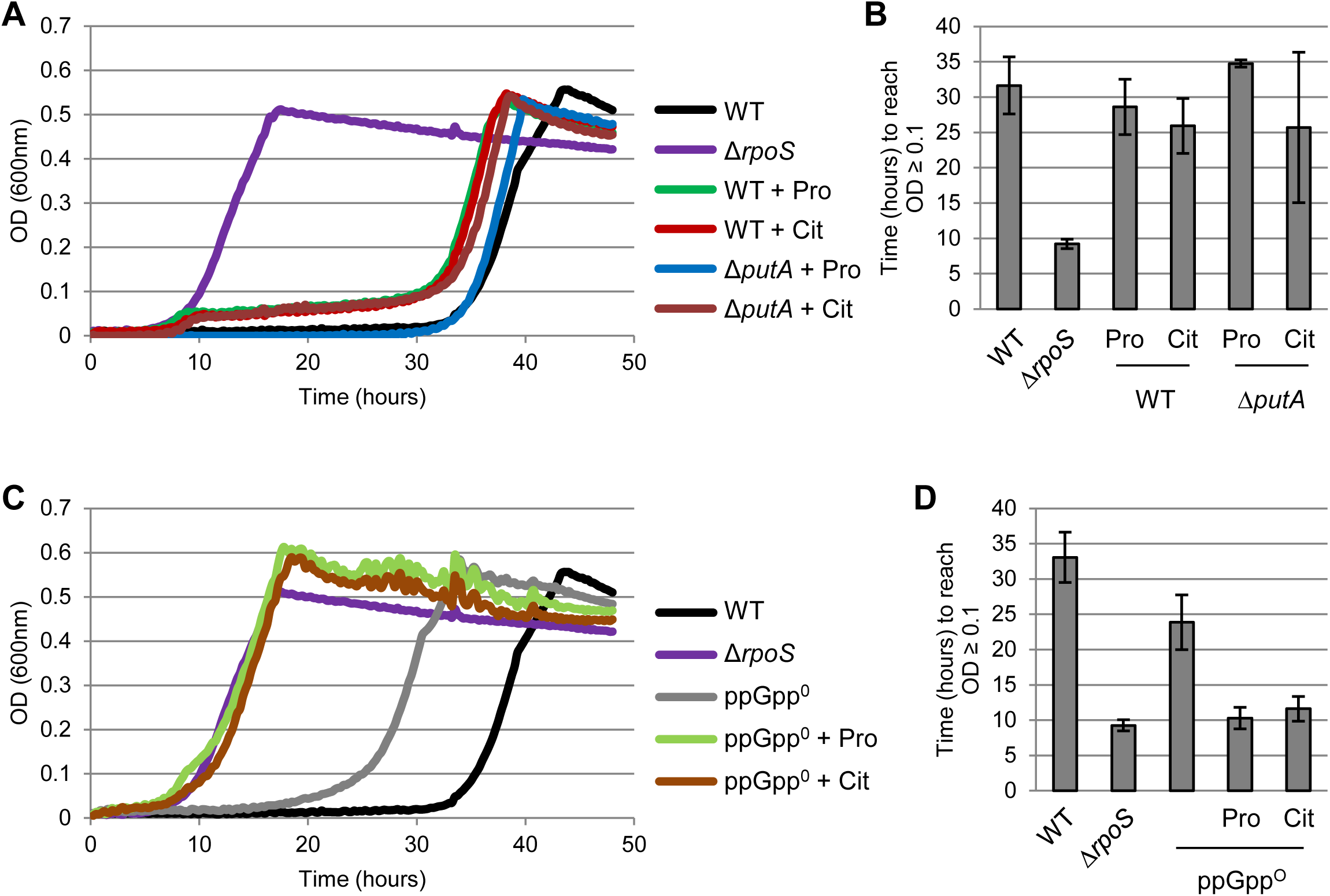
Specific nutrients induce growth using succinate and diauxic repression involves the stringent response. **A and C)** Representative growth curves of *Salmonella* growth in MOPS minimal media with 0.2% succinate as the primary carbon source and supplemented with 0.005% proline (+ Pro) or 0.005% citrate (+ Cit) where indicated. **B and D)** Graphs showing the average time in hours for *Salmonella* to reach an OD of 0.1 as an analog of the length of lag phase using succinate as the sole carbon source. The data shows the average of at least three biological replicates and error bars show one standard deviation.

We considered the possibility that proline or citrate are limiting nutrients during growth on dicarboxylates (e.g. proline synthesis is shut down and the cell becomes auxotrophic), however several additional observations suggest that this is not the case. We note, for example, that growth is temporarilty restored by either citrate or proline, and neither metabolite is known to be an intermediate in the synthesis of the other. Therefore, these two compounds are likely affecting growth through similar but independent routes.

Another observation relates to the effect of proline or citrate on the dicarboxylate growth lag in a *Salmonella relA/spoT* double mutant that is unable to produce the secondary messenger ppGpp to trigger the stringent response. We find that the *Salmonella* stringent mutant displays a growth lag similar to wild-type when subcultured into succinate media from an overnight culture grown in LB or M9 media. Whereas wild-type *Salmonella* rapidly shuts down growth upon exhaustion of available proline or citrate, the stringent mutant did not (Figure 4C and D) – that is the strain continued to grow on succinate.

This suggests that exhaustion of proline or citrate are not, in and of themselves, limiting during growth in succinate media, but rather that proline and citrate might act as external cues that allow growth on dicarboxylates to start. In wild-type *Salmonella* strains, growth continues until these stimulating compounds are exhausted; at which point the production of ppGpp via the stringent response stimulates the shutdown of dicarboxylate utilization, possibly through its effects on RpoS. In the absence of ppGpp or RpoS, however, the exhaustion of available proline/citrate is not sensed by the cell, and growth will continue until the dicarboxylate (or another essential nutrient) are exhausted.

### Proline-mediated stimulation in growth requires the multifunctional proline utilization enzyme PutA

Proline has been shown to play a key, but unclear, role in *Salmonella*’s adapation to life in an animal host. For both *E. coli* and *Salmonella*, proline can act as a compatible solute to regulate osmotic balance in the cells (34, 35). However, the key *Salmonella*-specific virulence factors encoded by *mgtA* and *mgtCBR* are controlled by short proline-rich peptides (*mgtL* and *mgtP*, respectively) that attenuate their operons in response to increased levels of uncharged tRNA_Pro_ (36). Also, loss of EF-P, which is critical to ensure the smooth translation of proteins rich in proline, causes translational stalling that likely mimics the effects of proline-limitation (30, 37, 38). Notably, *Salmonella* strains lacking EF-P are poorly virulent in animal models and appear to have difficulty adapting to an intraceulluar lifestyle; e.g. failing to produce stereotypical “sifs” when infecting host cells (29).

To explore the factors that might link proline availability to the rapid utilization of dicarboxylates as carbon sources, the roles of several proline-dependent regulatory systems were examined. While neither *mgtA* nor *mgtCBR* had an effect on proline’s effect on the dicarboxylate growth lag (data not shown), we found that strains lacking PutA, an unusual polyfunctional proline utilization enzyme, were unresponsive to proline with respect to growth on succinate (35, 39, 40).

PutA is a polyfunctional membrane-bound enzyme that contains the two enzymatic activities required to oxidatively convert proline to glutamate: proline dehydrongease (PRODH), which oxidizes proline to P5C; and P5C dehydrogenase (P5CDH) which is critical to convert P5C to glutamate. During the course of this conversion, PutA transfers resulting electrons to ubiquinone (via PRODH), and NAD+ (via P5CDH). PutA also contains a DNA binding domain that controls, via repression, its own expression (*putA*) as well as that of the divergently transcribed *putP* gene, encoding the proline importer PutP. An implication of this finding is that proline, *per se*, may not be the proximal metabolite that regulates the utilization of dicarboxylates, but instead may be due to PutA-mediated reduction of the quinone or NAD pool, the availability of gluatamine/glutamate (although neither glutamine nor glutamate had the same effect as proline) or PutA-mediated gene regulation via the PutA DNA-binding domain.

### Repression of dicarboxylate import accounts for growth lag

Activation of RpoS not only upregulates several systems directly involved in mitigating cellular stress (e.g. catalase, import and synthesis of compatible solutes, the production of glycogen, etc.), but it has also been shown to down regulate the expression of genes encoding enzymes in the TCA cycle (41–43). An analysis of *dctA* gene expression, encoding the primary dicarboxylate transporter, on SalCom (44, 45) showed that it generally anti-correlated with the expression of RpoS-activated genes like OsmY, where *dctA* was downregulated in response to osmotic shock and in late stationary phase. We hypothesized that activation of the RpoS-regulon may repress the expression of the *dctA* gene and thereby restrict *Salmonella* from taking up dicarboxylates such as succinate for consumption in response to stresses encountered within the host. To test if low *dctA* expression accounts for the *Salmonella* extended lag on dicarboxylates, we constitutively expressed *dctA* from a plasmid. Indeed, constitutive expression of *dctA* (but not of a similar *lacZ* control plasmid) eliminated the *Salmonella* lag in succinate media (Figure 5A and B).

**Figure 5:**
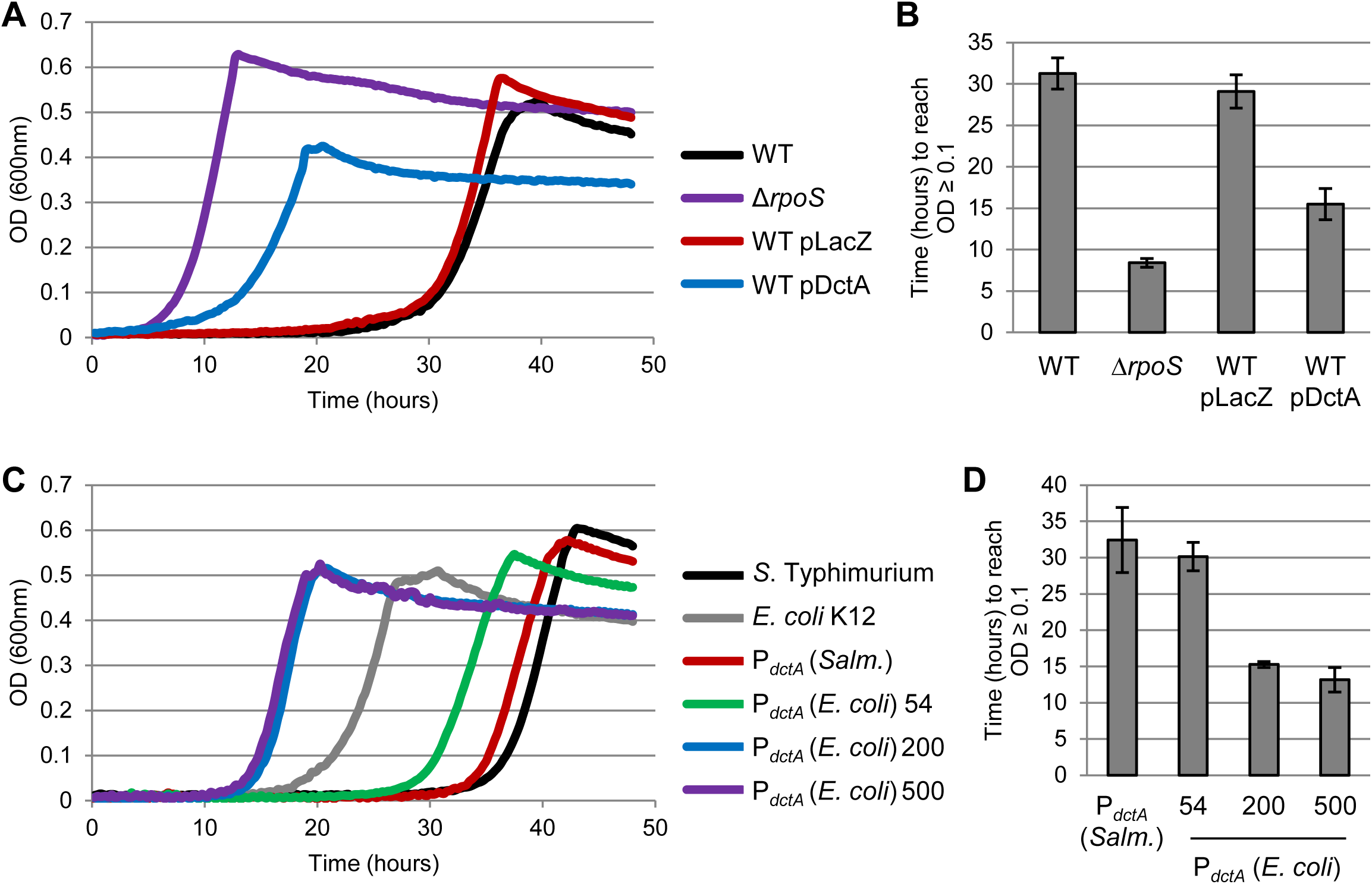
Expression of *dctA* induces growth using succinate. Growth of *Salmonella* in MOPS minimal media with 0.2% succinate as the sole carbon source. **A)** Overexpression of *dctA* from a plasmid. WT pLacZ and pDctA indicate wild-type *Salmonella* containing a pXG10sf plasmid encoding full length *lacZ* or *dctA* with expression driven by the constitutively active PLtet0-1 promoter. The *rpoS* mutant is shown for comparison. **B)** Graph showing the average time in hours for wild-type (WT) or mutant *Salmonella* to reach an OD of 0.1 as an analog of the length of lag phase using succinate as the sole carbon source. The data shows the average of at least three biological replicates and error bars show one standard deviation. **C)** Stretches of the *E. coli dctA* promoter (P_*dctA*_) were inserted into the *Salmonella* chromosome replacing the native *dctA* promoter. The length inserted (in base-pairs counting back from the *dctA* start codon) are indicated. Replacement with *Salmonella*’s native *dctA* promoter controls for effects due to the insertion method (P_*dctA*_ *Salm*.). Wild-type *Salmonella* and *E. coli* K12 are shown for comparison. **D)** As for B.

### The *E. coli dctA* promoter is sufficient to induce *Salmonella* growth using dicarboxylates

Given the differences between *Salmonella* and *E. coli*, with respect to growth on succinate, we compared the nucleotide sequences upstream of *dctA* in these bacteria (Figure S3). The two regions are largely similar (78% identity overall across 500bp upstream), especially near the core promoter. However, there are notable regions of significant sequence difference that are specific and conserved to each species. A comparison of the *dctA* promoter of *S. bongori* and of the various *S. enterica* subspecies (enterica, houtenae, arizonae, diarizonae, salmae) and typhoidal isolates reveal that they are highly conserved to one another - but distinct from the *dctA* promoters found in pathogenic and commensal *E. coli* strains. This suggests that changes to the regulation of *dctA* may have occurred as, or shortly after, the two species diverged.

To examine whether the species-specific differences in the *dctA* promoter (P_*dctA*_) played a role in the differences in how these species respond to succinate, the *E. coli* P_*dctA*_ was engineered into the *Salmonella* chromosome using an upstream chloramphenicol resistance cassette to select for successful recombination. Replacing the 500bp upstream of the *Salmonella dctA* start codon with those of *E. coli* was sufficient to abolish *Salmonella*’s ability to repress its uptake of dicarboxylic acids and this strain grew readily in succinate media (Figure 5C and D). As a control, using the same method to insert *Salmonella*’s native *dctA* promoter yielded no difference from wild-type *Salmonella*. Further reducing the swapped region to 200bp maintained the growth phenotype, but swapping only 54bp upstream of the *Salmonella* with that of *E. coli* (constituting the 5’ untranslated region) resulted in only a slight restoration of growth.

It is possible that *Salmonella* contains a factor to repress *dctA* expression that is not present in *E. coli*, so we generated transcriptional fusion plasmids and tested the two *dctA* promoters when expressed in *E. coli*. We found that the *dctA* promoter from *Salmonella* was expressed to a significantly lower degree than that of *E. coli* (Figure S4), suggesting that it contains a distinctive region that is recognized by a common regulator that is also present in *E. coli*.

### Restricting dicarboxylate import does not influence survival in macrophage cell lines

To probe the question of why *Salmonella* may have acquired the trait of blocking dicarboxylate utilization and what evolutionary advantage it may gain by it, we considered that succinate levels increase significantly in activated macrophage, an environment that *Salmonella* (but not *E. coli*) has adapted to survive in effectively (5, 6, 46). In the *Salmonella*-containing vacuole, the pH reaches approximately 5.0 and the lower estimates reach pH 4.4, which is comparable to the acid dissociation constants of succinate (pK_a1,2_ = 4.2, 5.6) (47, 48). This suggests that in the acidified phagosome, succinate may become protonated and potentially act as a proton shuttle to acidify the bacterial cytoplasm. The ability of *Salmonella* to restrict its uptake of succinate could therefore provide a possible survival advantage in this environment.

Constitutive overexpression of *dctA* but not *lacZ* led to decreased survival in both acidifed succinate media and in the human monocyte THP-1 cell line (Figure S5). However, overexpression of *dctA* has been demonstrated to be toxic to *E. coli* and we found that survival in acidified succinate was just as low for a *dctA* point mutant (N301A) that is defective for succinate transport (49). This implicated that the reduced survival was not due to dicarboxylate uptake but rather was an artifact of *dctA* overexpression to toxic levels. To bypass this artifact, we tested the *Salmonella* strain containing the chromosomal *dctA* promoter from *E. coli*, which grows readily in dicarboxylate media yet does not constitutively overexpress *dctA* from a plasmid and so does not exhibit the associated toxic effects. Using this strain we found no decrease in survival relative to wild-type *Salmonella* in acidifed succinate media or in human (THP-1) or mouse (J774) macrophage cell lines (Figure 6). Deletion of *dctA* or *iraP* genes also did not appear to significantly influence *Salmonella* survival in THP-1 macrophage in these short-term infection assays.

**Figure 6:**
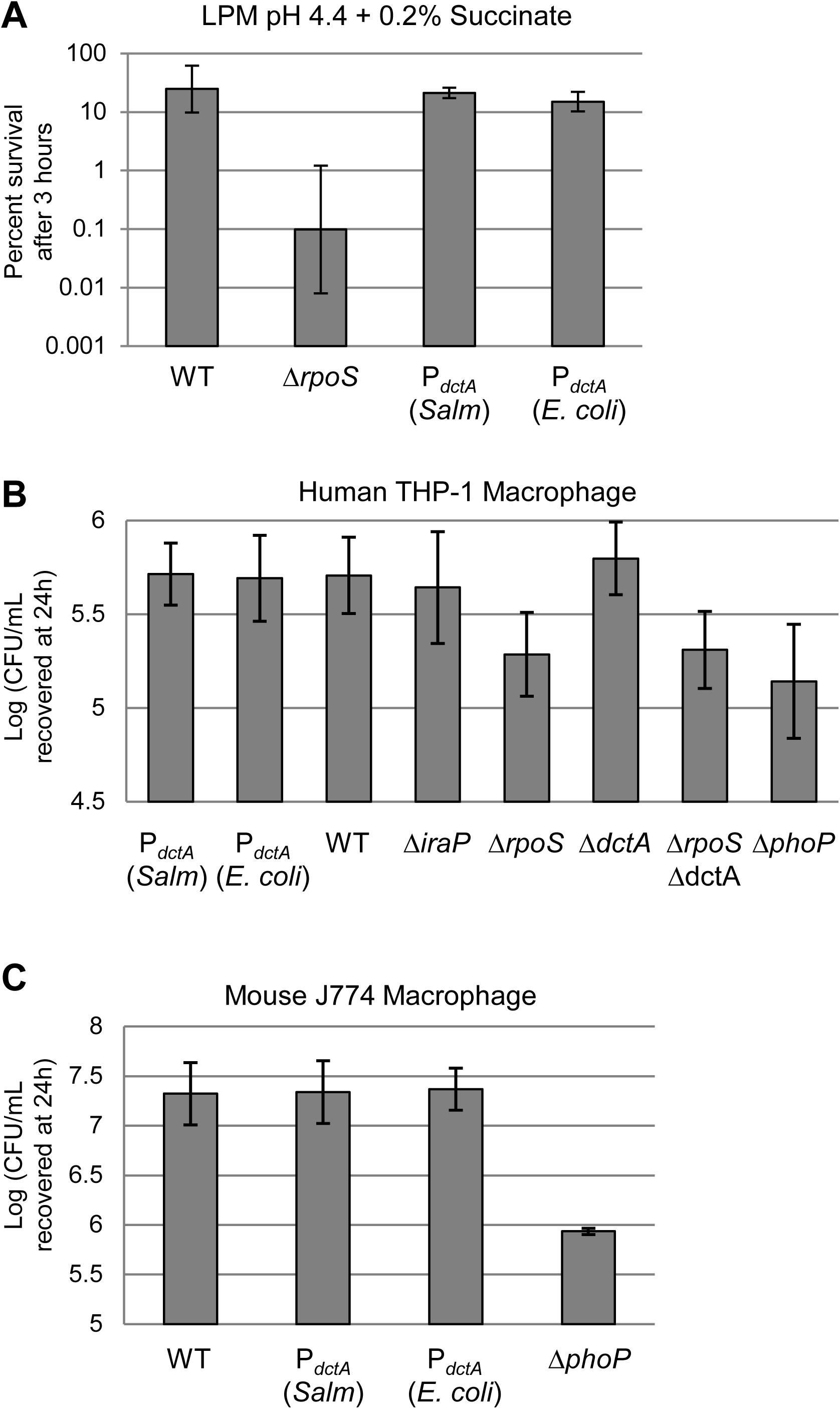
Replacement of the *Salmonella dctA* promoter does not influence survival in acidified succinate or macrophage. **A)** Survival of *Salmonella* treated with LPM media containing 0.2% succinate and acidified to pH 4.4. Wild-type (WT) is compared to *Salmonella* with its chromosomal *dctA* promoter replaced with 500bp of the *E. coli dctA* promoter (P_*dctA*_ *E. coli*) or *Salmonella*’s native *dctA* promoter as a control (P_*dctA*_ *Salm*). An *rpoS* mutant is shown as a positive control. CFU recovered at 3 hours were normalized to input CFU at 0h and expressed as percent survival. Data shows the average across three biological replicates and error bars indicate one standard deviation. **B)** Infection of THP-1 human macrophage comparing survival of wild-type *Salmonella* (WT) with various mutant strains and the *dctA* promoter swap strain. A *phoP* mutant is included as a positive control. Data shows a logarithm of CFU recovered at 24 hours post infection and is the average of three biological replicates. Error bars show one standard deviation. **C)** As in (B) but showing CFU recovered from mouse J774 macrophage.

## Discussion

In this work we explore a previously uncharacterized *Salmonella* phenotype whereby it restricts its ability to import dicarboxylates in an RpoS and Elongation Factor P dependent manner. Stationary phase cells subcultured into media with succinate, fumarate, or malate as the sole carbon source fail to grow for >30 hours before resuming normal growth. The cause of this extended lag phase appears to largely reflect low expression of the *dctA* gene encoding a major dicarboxylate importer. The ‘cue’ that induces cells to suddenly begin rapid growth on dicarboxylates after 30 hours remains unclear. Interestingly, an almost immediate exit from the extended lag phase could be triggered by the addition of low amounts of proline or citrate. Upon exhaustion of the available proline or citrate, growth would again cease in a process that required the synthesis of ppGpp.

Our model suggests that activation of RpoS leads to a rapid shutdown in the transporter that mediates the uptake and utilization of dicarboxylates (Figure 7). Since the stringent response and ppGpp can impact the expression of *rpoS* and RssB anti-adaptors including *iraP*, the stringent response is likely the means by which IraP- and RpoS-mediate the shutdown of growth (14, 50–52). We also note the many curious links that have been found between regulation of genes essential for *Salmonella* host infection and the availability of proline, notably the attenuation peptides MgtL and MgtP and the essentiality of EF-P to alleviate translational stalling at proline rich motifs. Our finding that the poly-functional proline metabolic enzyme, PutA, is essential for the proline-mediated induction of growth on succinate, leaves open the question of whether the inducing cue is due to changes in the abilty of PutA to bind DNA (perhaps PutA directly represses *dctA* in Salmonella), alterations in cellular energetics via the production of either reduced quinones or NADH, or through increases in the concentrations of cellular glutamate (although we note addition of glutamate does not have the same effect as proline).

**Figure 7:**
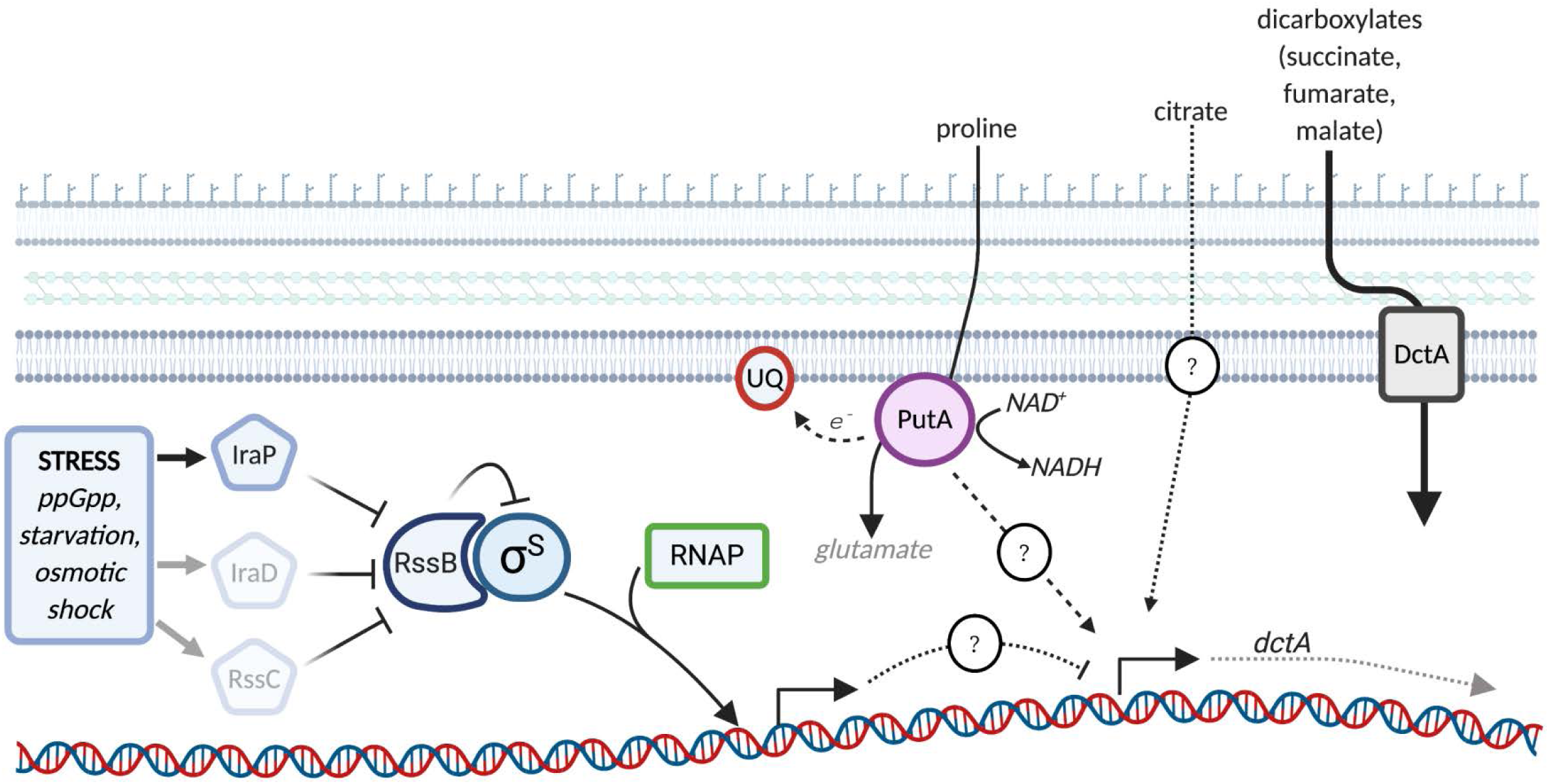
Model of the link between stress and dicarboxylate utilization in *Salmonella*. Under conditions of stress RpoS (σ^S^) levels are elevated due to the production of anti-adaptors that block the degradation of RpoS by the RssB adaptor and ClpXP protease system. RpoS directs RNA polymerase (RNAP) to several promoters involved in stress protection as well as to a series of downstream regulatory genes (including small RNAs) that decrease the production of enzymes involved in the TCA cycle and other metabolic pathways. One of these downstream regulatory factors, yet to be identified, impinges on the expression of *dctA*. The presence of either citrate or proline is sufficient to alleviate the block on dicarboxylate utilization. The proline mediated effect requires PutA, a polyfunctional enzyme that converts proline to glutamate with the concomitant production of reducing equivalents that are transferred to NAD+ and ubiquinone (figure created with BioRender). The factors that mediate the citrate-dependent effect remain unclear.

The difference in the response of *Salmonella* and *E. coli* to dicarboxylic acids may offer important clues to identifying the evolutionary advantage conveyed by this adaptation. The divergent evolutionary paths between *Escherichia* and *Salmonella*, that began over 100MYr ago, were fueled both by the acquisition of novel functions via horizontal gene transfer and extensive regulatory re-wiring, often through changes in cis-regulatory elements (53–56). The differences in the severity of their dicarboxylate lag, as evidenced by sampling a range of isolates in the SGSC and ECOR collections, appears to be due to extensive differences in the 200 nucleotide region upstream of the *dctA* dicarboxylate import gene including regions that lie upstream of the previously mapped 5’ UTR. While it remains possible that this factor could involve a small RNA, our finding that swapping just the 5’ untranslated region of *dctA* (P_*dctA*_ 54) was insufficient to reverse growth repression suggests a protein factor acting on the promoter at the transcriptional level. RpoS likely impacts *dctA* expression via an intermediate and yet undetermined transcription factor.

Since many of the traits that *Salmonella* has acquired since their divergence are related to its pathogenic lifestyle it follows that this phenotype may reflect a situation that *Salmonella* encounters during infection of a host. Interestingly, it has recently been shown that *Salmonella* utilizes microbiota-derived succinate in the lumen of the inflamed gut (57). This suggests that the uptake of succinate in the gut employs anaerobic dicarboxylate transporters rather than DctA, or that an inducing compound such as proline or citrate is present in this environment and can stimulate succinate uptake. The recent finding that succinate accumulates to high levels in activated macrophage suggests that *Salmonella*’s intracellular survival may represent the crucial selective environment that has led to the *dctA*-repression phenotype (46). It is conceivable that *Salmonella* recognizes the succinate produced by activated macrophage and restricts its uptake of dicarboxylates in response. Our examination of *Salmonella* strains that are impaired in their regulation of dicarboxylate uptake identified no survival defect in acidifed succinate or in macrophage cell lines. However, it remains possible that this repression phenotype is related to other aspects of the *Salmonella* virulence cycle beyond short-term survival in phagocytes. For instance, *Salmonella* may potentially restrict its import and consumption of dicarboxylates to maximize macrophage succinate levels leading to the production of the pro-inflammatory citokine IL-1β (46). This would not grant *Salmonella* an advantage for survival in macrophage *per se*, but would give *Salmonella* remaining in the gut lumen a growth advantage by maximizing the immune system’s oxidative burst and subsequent production of tetrathionate, a compound that *Salmonella* is distinctively equipped to use as a terminal electron acceptor (58– 60). Thus, if *Salmonella* were to import and consume succinate in macrophage, the inflammatory response might be deterred, restricting the ability of *Salmonella* to outcompete the gut microbtioa during infection.

We did note a few *Salmonella* strains that grew early on succinate despite having a catalase positive (and presumably *rpoS*+) phenotype. In all instances where multiple strains from the same *Salmonella enterica* serovar were tested, at least one exhibited an extended lag phase (Table S1). Thus, the earlier growth does not appear to be a trait of particular serovars but rather may reflect individual strains having lost (or never acquired) the delayed growth phenotype. This could occur by mutations in genes other than *rpoS*, such as *efp* or *iraP*, that permit early growth in dicarboxylic acids. Other exceptions include multiple *E. coli* strains that show an extended lag in their growth using succinate. While it is possible that these have mutations in genes required for the uptake of succinate (such as *dctA* or *dcuSR*), these may reflect genuine variation in how *E. coli* strains respond to dicarboxylates.

Finally, our findings have important implications for the interpretation of previous metabolic studies in commonly used laboratory strains. For example, it is important to consider that the Biolog assays primarily use succinate as the carbon source under many conditions (28, 29, 31). Thus, what we previously assumed to be improved growth of the *efp* mutant in a variety of different conditions in fact resulted solely from its ability to rapidly utilize dicarboxylate media. We also note the large number of metabolic studies carried out with the commonly used *Salmonella* Typhimurium strain, LT2, which is a known *rpoS* mutant (32). Because of this, studies using the LT2 strain would behave quite differently in Biolog assays than the other commonly used ‘wild-type’ *Salmonella* strains SL1344 (of which many isolates are auxotrophic for histidine) and 14028s.

## Materials and Methods

### Bacterial strains and plasmids

Bacterial strains, plasmids and primers are listed in Supplementary Tables S2-S4. As described previously, lambda red recombination (28, 61) and subsequent P22 phage transduction (62) was used to generate all of the gene knockout mutants in *Salmonella enterica subsp. enterica* serovar Typhimurium (*S*. Typhimurium) strain 14028s. *E. coli* was from the Keio collection K12 BW25113 strain background (63). To sample the genetic diversity of *Salmonella* and *E. coli* isolates, the *Salmonella* genetic stock centre (SGSC) SARA (64), SARB (65), and SARC (66) collections were employed and compared to the *E. coli* reference (ECOR) collection (67).

The full length *DctA* ORF was expressed from the pXG10sf plasmid under the control of the constitutively active PLtet0-1 promoter (68, 69). The IraP complementation plasmid was generated by inserting the *iraP* ORF and the upstream 300bp into pXG10sf. For promoter expression, the *dctA* promoter (500bp upstream of the *dctA* start codon) was inserted into pXG10sf to drive expression of superfolder GFP (69, 70). To generate the chromosomal *dctA* promoter swap strain, 500bp upstream of the *E. coli dctA* start codon, along with a chloramphenicol resistance cassette for selection, was inserted into the corresponding location of the *Salmonella* chromosome.

### Growth using dicarboxylates as a sole carbon source

Overnight LB cultures inoculated from single colonies were resuspended in MOPS minimal media with no carbon source to an optical density (OD_600_) of approximately 1.75. This suspension was used to inoculate (1/200 dilution) MOPS minimal media containing 0.2% carbon source (succinate unless otherwise indicated). Growth was conducted in a TECAN Infinite M200 plate reader at 37°C with shaking and OD_600_ was read every 15 minutes. For salts and hydrates of carbon sources the final concentration reflects the percent of the carbon source itself (e.g., 0.2% succinate was made as 0.47% sodium succinate dibasic hexahydrate).

For the SGSC and ECOR collections screen, 47 strains were assessed in duplicate per run in a 96-well plate. Wild-type and *rpoS* mutant *Salmonella* were included on every plate as quality controls. Each strain was tested on at least three separate days.

### Catalase assay

For each replicate of the SGSC and ECOR collections screen, each strain was tested for catalase activity as an analog for RpoS function (71). In parallel to the LB overnight cultures used as inoculum, 10μl of each culture was spotted onto an LB plate. The next day the spots were tested for catalase activity by the addition of 10μl hydrogen peroxide. Bubbling was scored compared to wild-type (catalase positive) and *rpoS* mutant (catalase negative) *Salmonella*.

### Acid survival

LPM media was made as described previously (72) and succinate was added to either 0.2% or 0.4% as indicated in figures. The pH of the media was then adjusted to 4.4. LB overnight cultures were resuspended to an OD of 0.1 in acidified media and incubated in a 37°C water bath. At time points, samples were taken, serial diluted and plated for colony forming units (CFU).

### Intra-macrophage survival

The THP-1 human monocyte cell line and the J774 mouse macrophage cell line were maintained in RPMI Medium 1640 (with L-glutamine) supplemented with 10% FBS and 1% Glutamax, and grown at 37°C and 5% CO_2_. For infection assays, THP-1 cells were seeded in 96-well plates at 50,000 per well with 50nM PMA (phorbol 12-myristate 13-acetate) added to the media to induce differentiation to adherent macrophage. After 48h, the media was replaced with normal growth media (no PMA) overnight. For infections with J774 macrophage the cells were seeded in 96-well plates at 50,000 per well overnight. *Salmonella* in RPMI were added onto seeded cells at a multiplicity of infection (MOI) of approximately 20 bacteria to 1 macrophage and centrifuged for 10 minutes at 1000rpm for maximum cell contact. After centrifuging the plate was placed at 37°C (5% CO_2_) and this was called ‘time 0’. After 30 minutes, non-adherent *Salmonella* were washed off by three washes with PBS followed by replacement with fresh media containing 100 μg/ml gentamicin to kill extracellular *Salmonella*. At 2 hours the media was replaced with media containing gentamicin at 10 μg/ml. At timepoints, intracellular bacteria were recovered using PBS containing 1% Triton X-100 and vigorous pipetting. Samples were serially diluted and five 10μl spots were plated for CFU counting. Each sample included three separate wells as technical replicates (a total of 15 x 10μl spots counted per biological replicate).

## Acknowledgements

We would sincerely like to thank Dr. Scott Gray-Owen and members of his lab, in particular Dr. Ryan Gaudet, for their generous donation of technical expertise, macrophage cells lines, and use of their equipment. WWN was supported by an Operating Grant from the Canada Institutes for Health Research (MOP-86683) and a Natural Sciences and Engineering Research Council (NSERC) of Canada Grant (RGPIN 386286-10). SJH was supported by an NSERC Vanier Canada Graduate Scholarship.

## Supplemental Materials and Methods

### Promoter induction fluorescence

Growth was conducted in a TECAN Infinite M200 plate reader at 37°C with shaking and OD_600_ and GFP fluorescence (475nm and 511nm excitation and emission wavelengths respectively) were read every 15 minutes. For clarity, bar graphs show fluorescence at 16h post inoculation. Chloramphenicol was included in all media at a concentration of 20μg/ml to maintain the plasmids.

## Supplemental Figures

**Figure S1:**
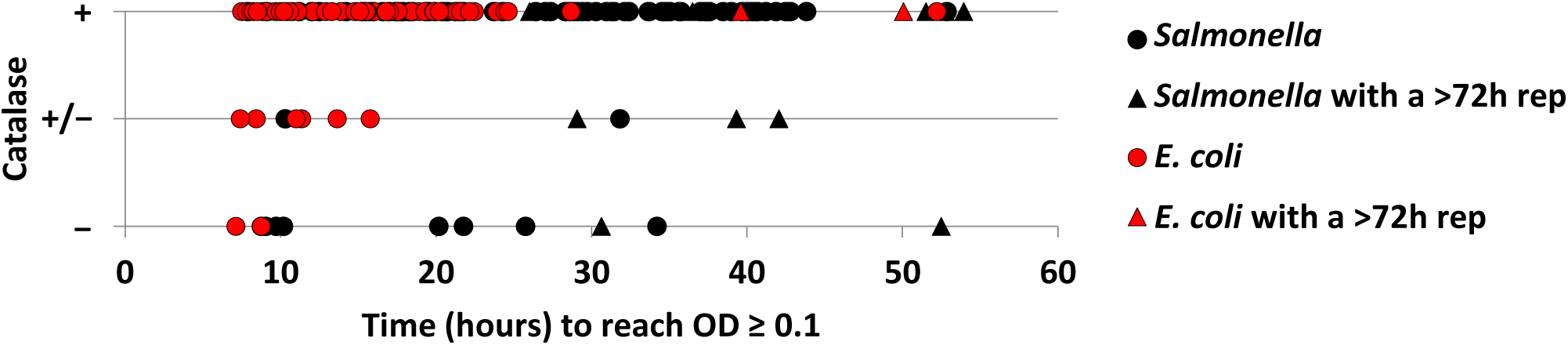
SGSC *Salmonella* trend towards longer lag phase than ECOR *E. coli* regardless of catalase activity. Comparing time to reach OD 0.1 in succinate media with catalase result for all strains tested from the SGSC and ECOR collections. + or − indicate that the strain showed positive or negative (respectively) catalase result in all replicates. +/− indicates either delayed positive or inconsistent catalase results across replicates. Data is the average of at least three biological replicates with the exception of strains shown as triangles that had at least one replicate that did not reach OD 0.1 by 72h (shown as average of the remaining replicates).

**Figure S2:**
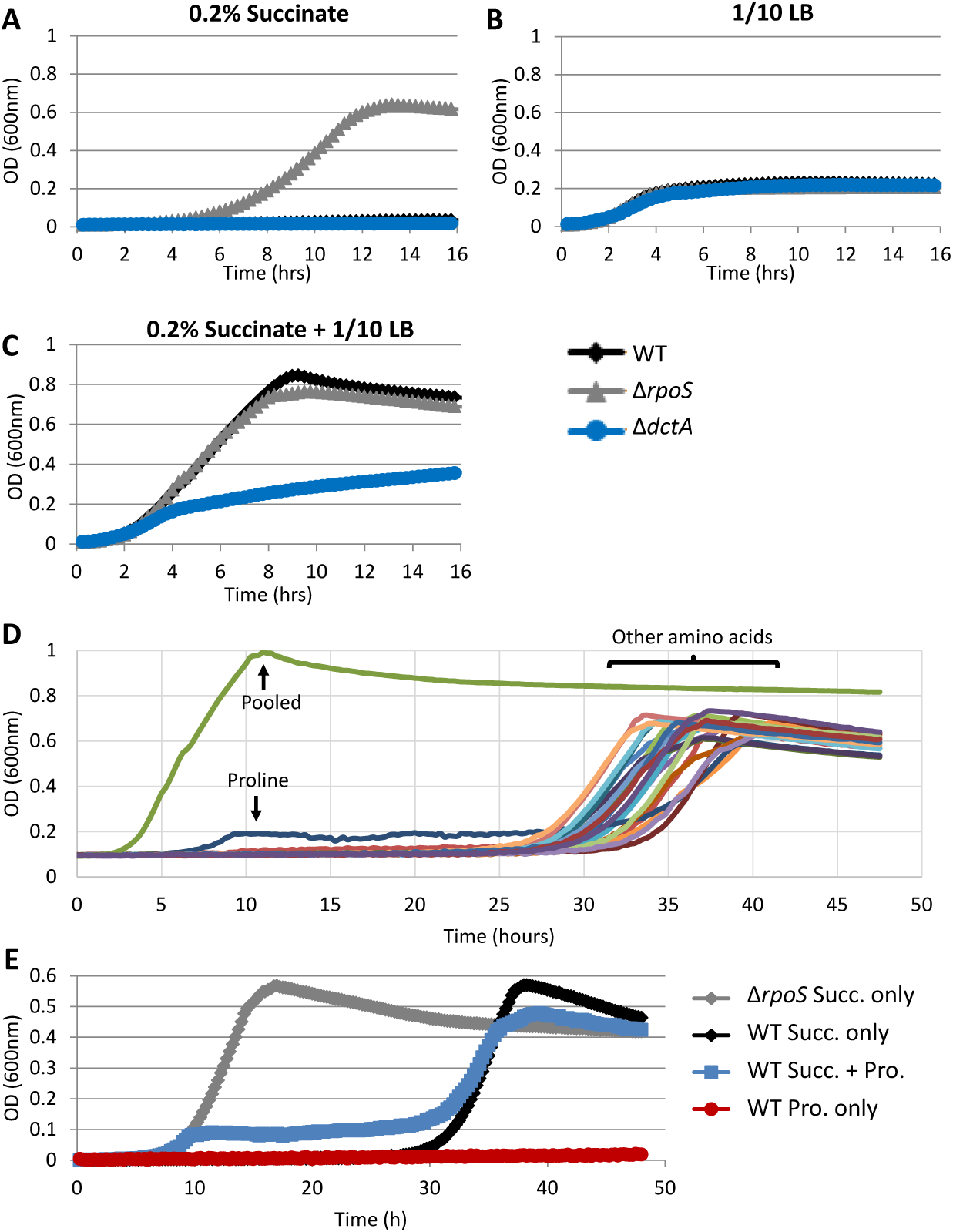
Growth in succinate media is induced by dilute amino acids. Growth of *Salmonella* in MOPS minimal media using **A)** 0.2% succinate as the sole carbon source, **B)** 1/10 diluted LB as the sole source of carbon, **C)**, both 0.2% succinate and 1/10 diluted LB. Legend for A-C is indicated at right. **D)** Growth of wild-type *Salmonella* in MOPS minimal media with 0.2% succinate as the primary carbon source and supplemented with individual or pooled amino acids at a concentration approximately 1/10 of that in LB media (1). **E)** Growth of *Salmonella* wild-type (WT) or *rpoS* mutant (Δ*rpoS*) in MOPS minimal media with 0.2% succinate (Succ.) and/or 0.01% proline (Pro.) as the sole carbon sources. Graphs are representative of at least two independent replicates.

**Figure S3:**
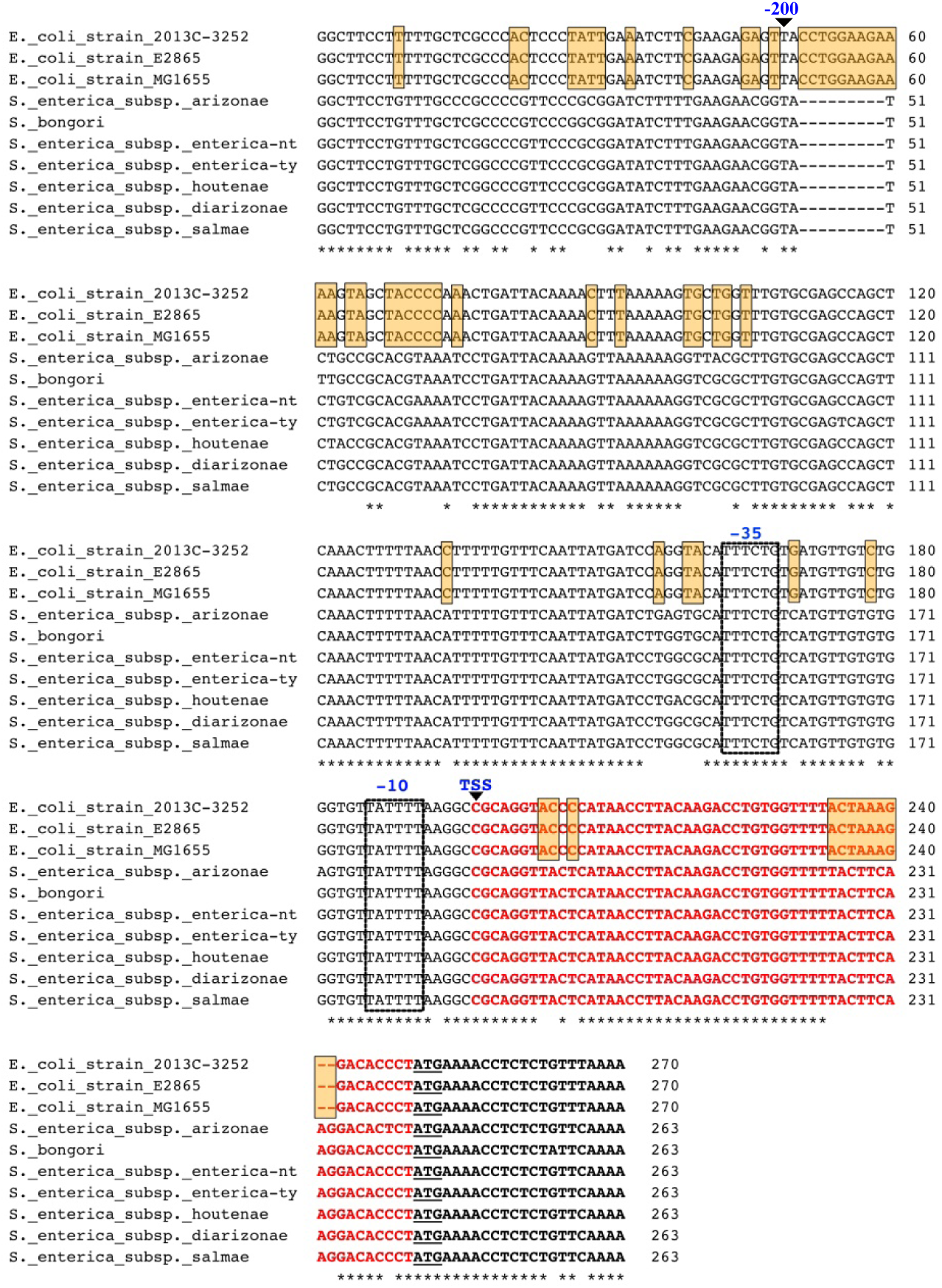
**Alignment of *Salmonella* and *E. coli dctA* promoters** conducted using Clustal Ω software (2). Asterisks indicate identity and regions of consistent differences between *E. coli* and *Salmonella* are highlighted in yellow. The 200bp from *E. coli* that were swapped into the *Salmonella* chromosome in Figure 5C and D are indicated (−200), as are the predicted −35 and −10 promoter elements identified by RegulonDB (3) and supplemental reference (4). The transcriptional start site (TSS) is indicated in blue with red text indicating the 5’ UTR as identified by RegulonDB (3) and in references (4, 5). The ATG start codons are underlined.

**Figure S4:**
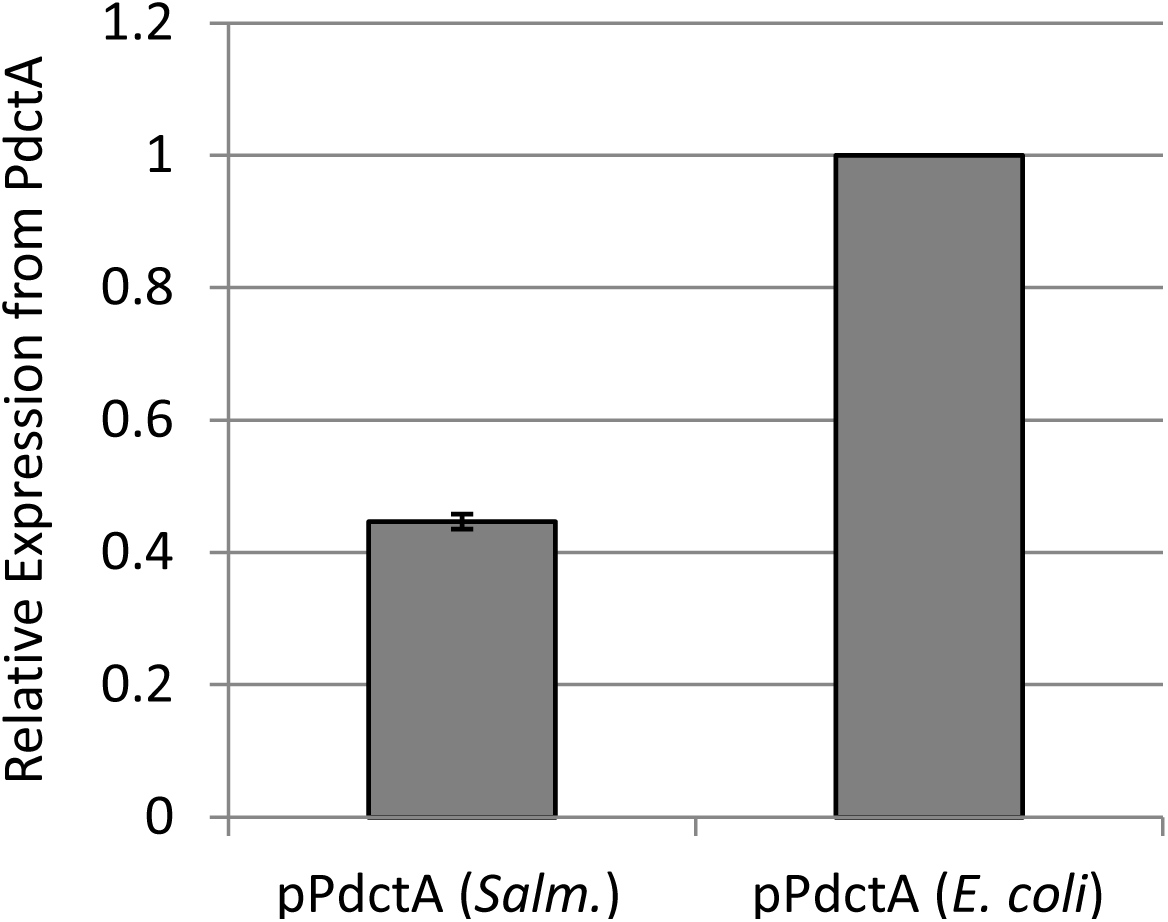
The *Salmonella dctA* promoter shows reduced expression in *E. coli* relative to the *E. coli dctA* promoter. Trancriptional fusions to GFP on a plasmid of the *Salmonella* or *E. coli dctA* promoters. GFP fluorescence was examined in *E. coli* K12 and data shows relative expression at 48h post-inoculation into MOPS minimal media with 0.2% succinate as the sole carbon source. Data is the average of three biological replicates and error bars show one standard deviation.

**Figure S5:**
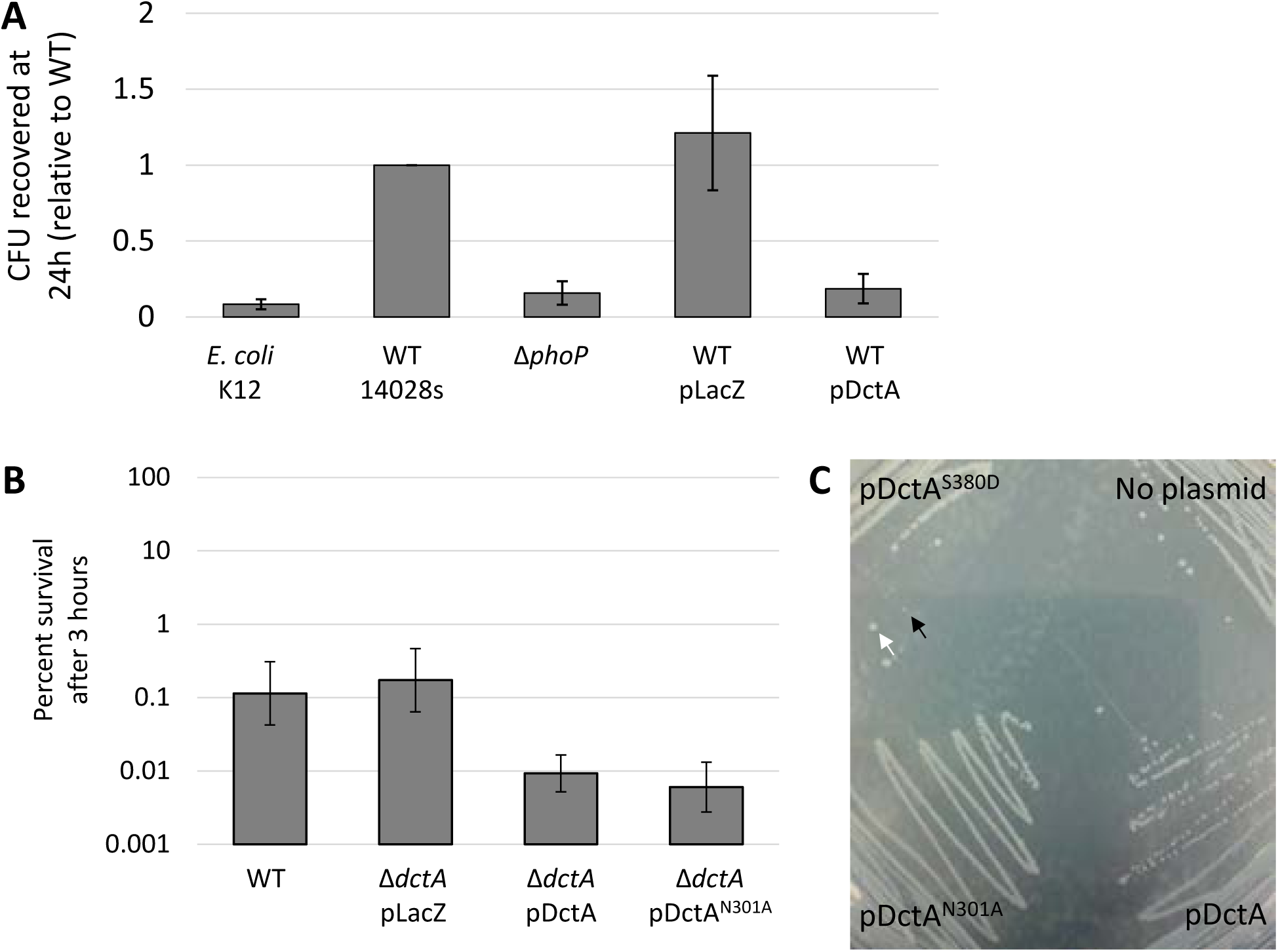
Overexpression of DctA from a plasmid decreases survival in macrophage due to toxicity. **A)** Infection of THP-1 macrophage examining survival of wild-type *Salmonella* with no plasmid (WT) compared to isogenic strains constitutively expressing LacZ (pLacZ) or DctA (pDctA) from the pXG10sf plasmid. Survival of *E. coli* and Δ*phoP* mutant *Salmonella* are included as controls for comparison. Data shows CFU recovered at 24 hours post-infection relative to wild-type *Salmonella* and is the average of three biological replicates. Error bars show one standard deviation. **B)** Survival of wild-type or *dctA* mutant *Salmonella* treated with LPM media containing 0.4% succinate and acidified to pH 4.4. CFU recovered at 3 hours were normalized to input CFU at 0h and expressed as percent survival. Data shows the average across three biological replicates and error bars indicate one standard deviation. Where indicated, genes were expressed from the ampicillin resistant version of the pXG10sf plasmid under the control of the constitutively active PLtet0-1 promoter. **C)** Representative image showing overnight growth of wild-type *Salmonella* on an LB agar plate comparing colony size when constitutively expressing *dctA* point mutants. Black arrow indicates a small colony of the strain expressing DctA^S380D^; white arrow indicates a large colony likely resulting from a suppressor mutation.

**Table S2:** Strains*

| Strain | Genotype | ID | Description | Reference |
| --- | --- | --- | --- | --- |
| *Salmonella*<br>Typhimurium<br>14028S |  | WN150 | Wild-type *Salmonella enterica* serovar Typhimurium strain 14028S | ATCC |
| | $\Delta efp$ | WN1405 | Deletion of *efp* gene in WN150 background. Resistance cassette was been removed by FLP recombination | (6) |
| | $\Delta rpoS::kan-FRT$ | WN2881 | Deletion of *rpoS* gene in WN150 background | This work |
| | $\Delta dctA$ | WN2914 | Deletion of *dctA* gene in WN150 background. Resistance cassette was been removed by FLP recombination | This work |
| | $\Delta iraP$ | WN3186 | Deletion of *iraP* gene in WN150 background. Resistance cassette was been removed by FLP recombination | This work |
| | $\Delta rssB::cm-FRT$ | WN3162 | Deletion of *rssB* gene in WN150 background | This work |
| | $\Delta rssC::cm-FRT$ | WN3163 | Deletion of *rssC* gene in WN150 background | This work |
| | $\Delta iraD::cm-FRT$ | WN3165 | Deletion of *iraD* gene in WN150 background | This work |
| | $\Delta putA::kan-FRT$ | WN2963 | Deletion of *putA* gene in WN150 background | This work |
| | $\Delta relA::cm-FRT$<br>$\Delta spoT30$ (*sms-211*)::*Tn10* | WN138 | Deletion of *relA* and *spoT* genes in 14028S background. Called ppGpp<sup>0</sup> in text and figures | This work |
| | $\Delta phoP::cm-FRT$ | WN152 | Deletion of *phoP* gene in 14028S background. Also known as CS015 | (7) |
| | $P_{dctA}(Salm.)$ | WN3188 | *Salmonella* *dctA* promoter with upstream chloramphenicol resistance cassette in the chromosome of the WM150 background. | This work |
| | $P_{dctA}(E. coli) 500$ | WN3189 | *E. coli* *dctA* promoter (500bp upstream of the start codon) with upstream chloramphenicol resistance cassette in the chromosome of the WM150 background. | This work |
| | $P_{dctA}(E. coli) 200$ | WN3239 | *E. coli* *dctA* promoter (200bp upstream of the start codon) in the chromosome of the WM150 background. -500 to -200bp remain from the *Salmonella* promoter preceded by an upstream chloramphenicol resistance cassette. | This work |
| | $P_{dctA}(E. coli) 54$ | WN3238 | *E. coli* *dctA* 5' UTR in the chromosome of the WM150 background. -500 to -54bp remain from the *Salmonella* promoter preceded by an upstream chloramphenicol resistance cassette. | This work |
| *E. coli* K12 BW25113 |  | WN0970 | Wild-type *E. coli* strain K12 | (8) |

\* Additional strains from the SGSC and ECOR collections are listed in Table S1

**Table S3:**
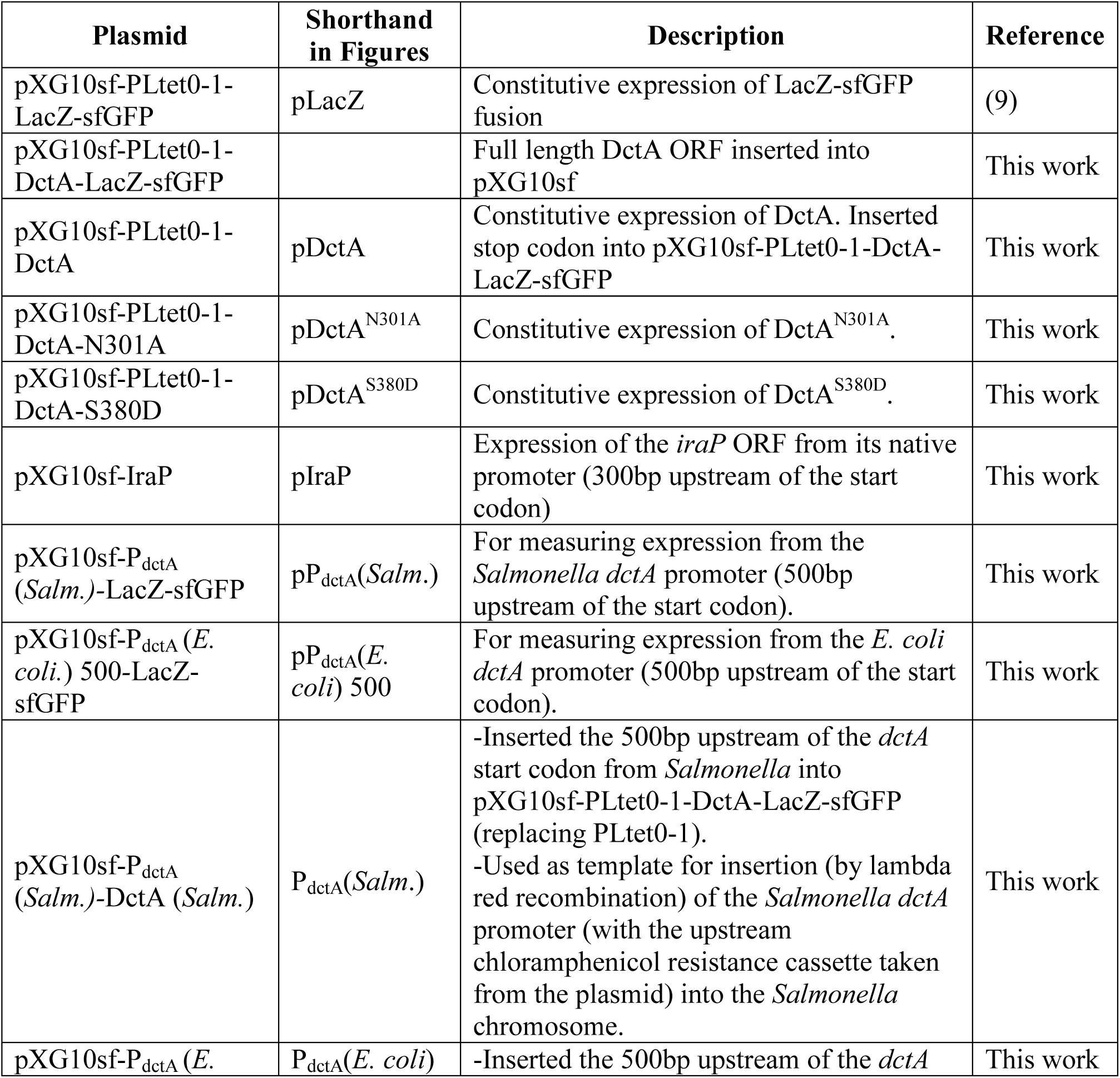
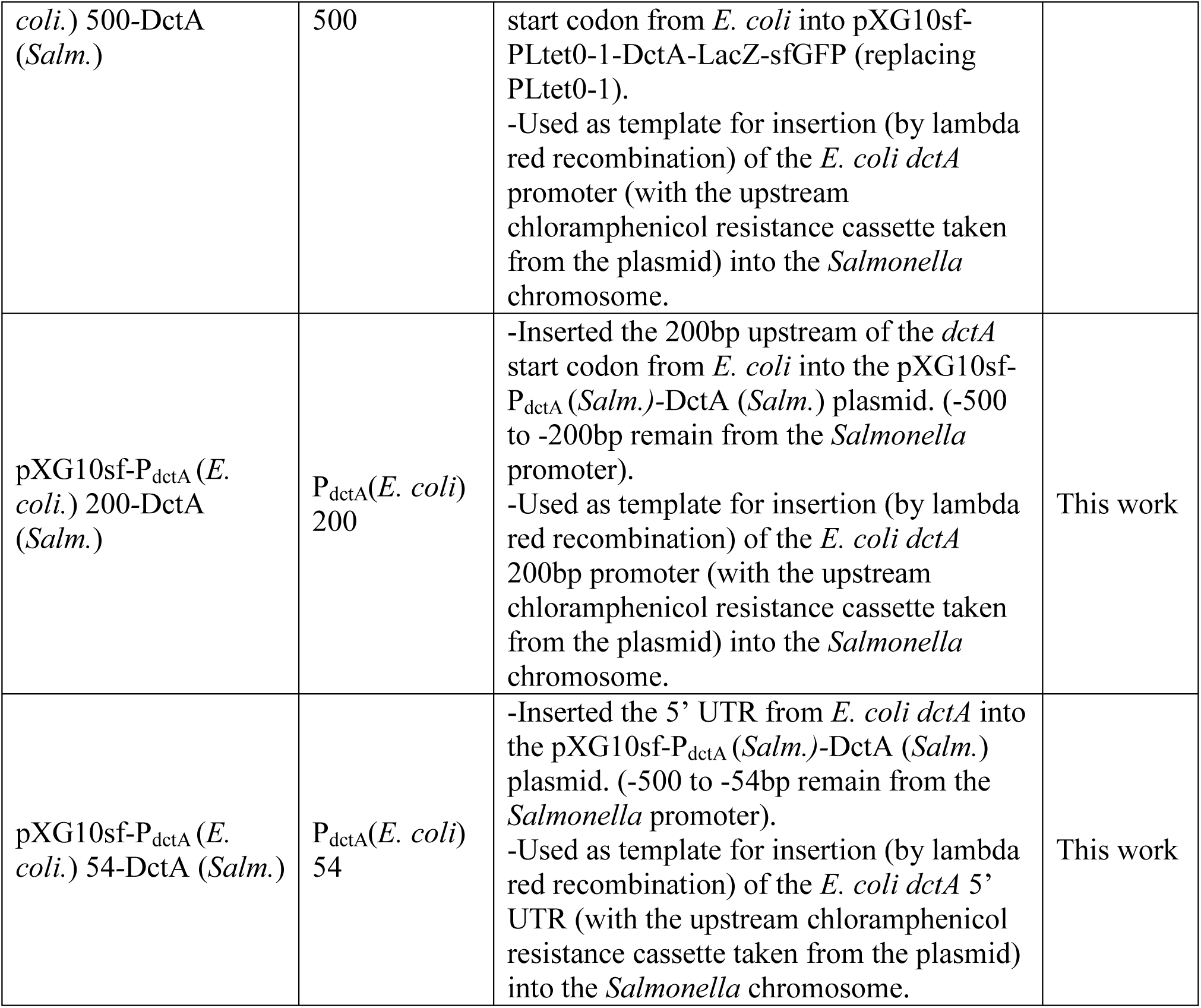
Plasmids

**Table S4:**
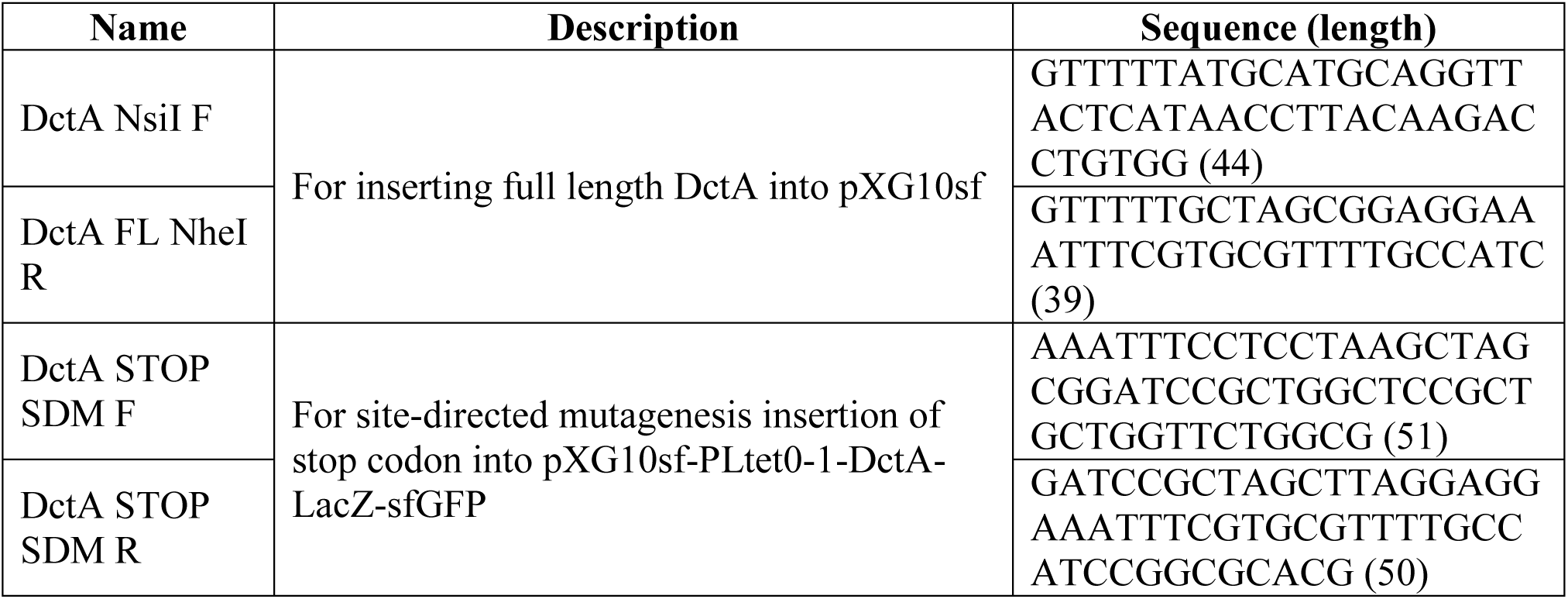
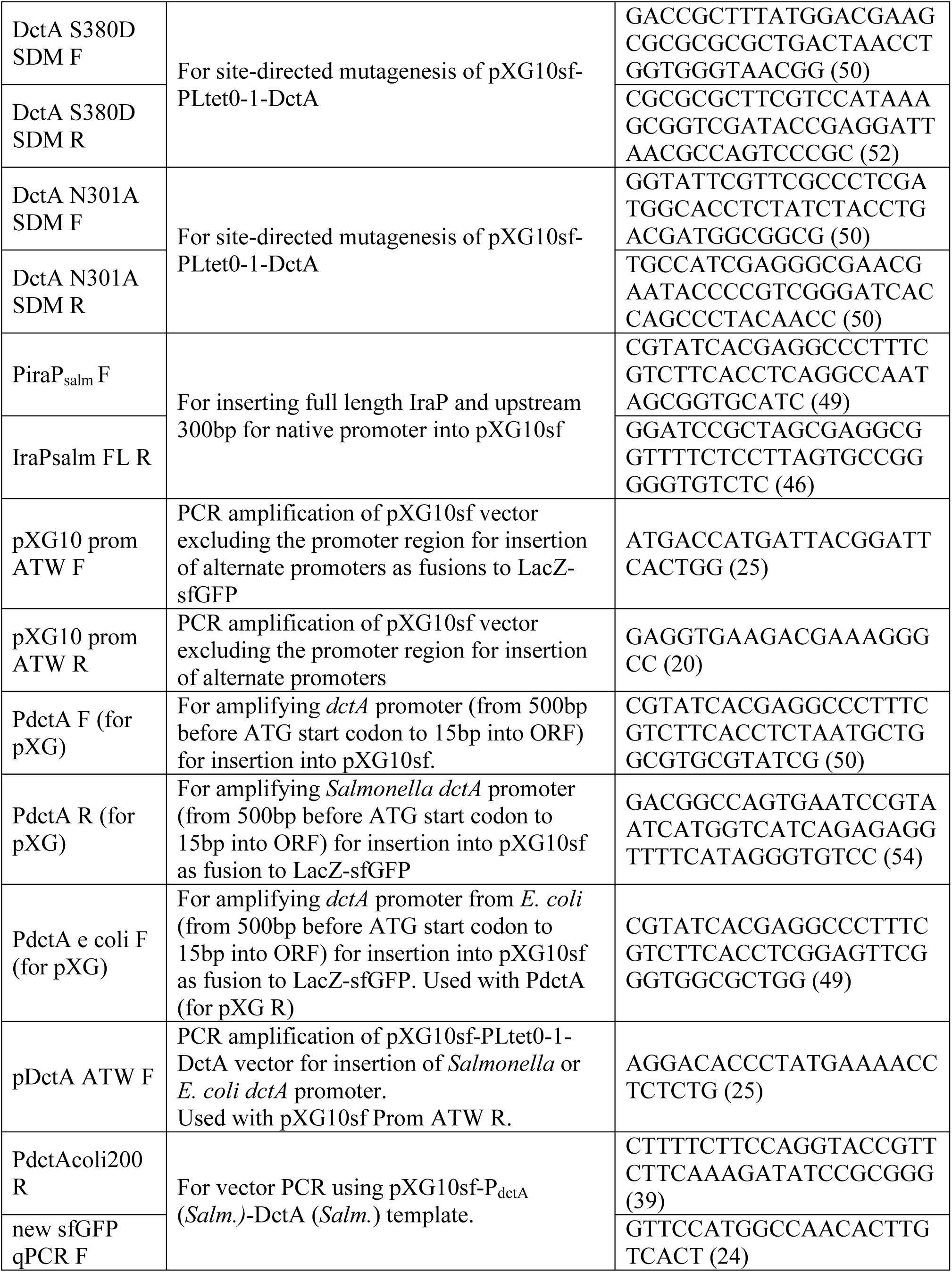
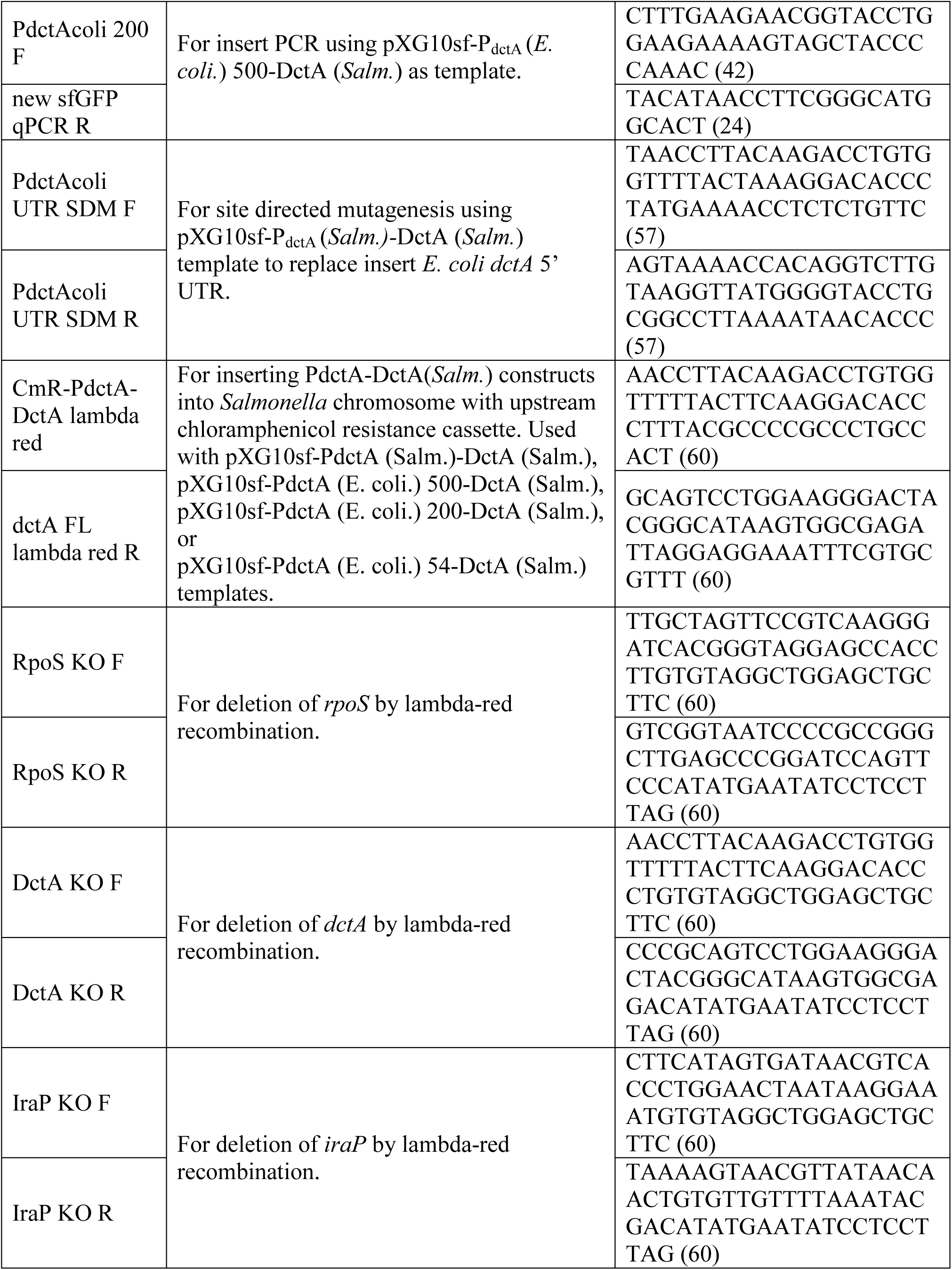
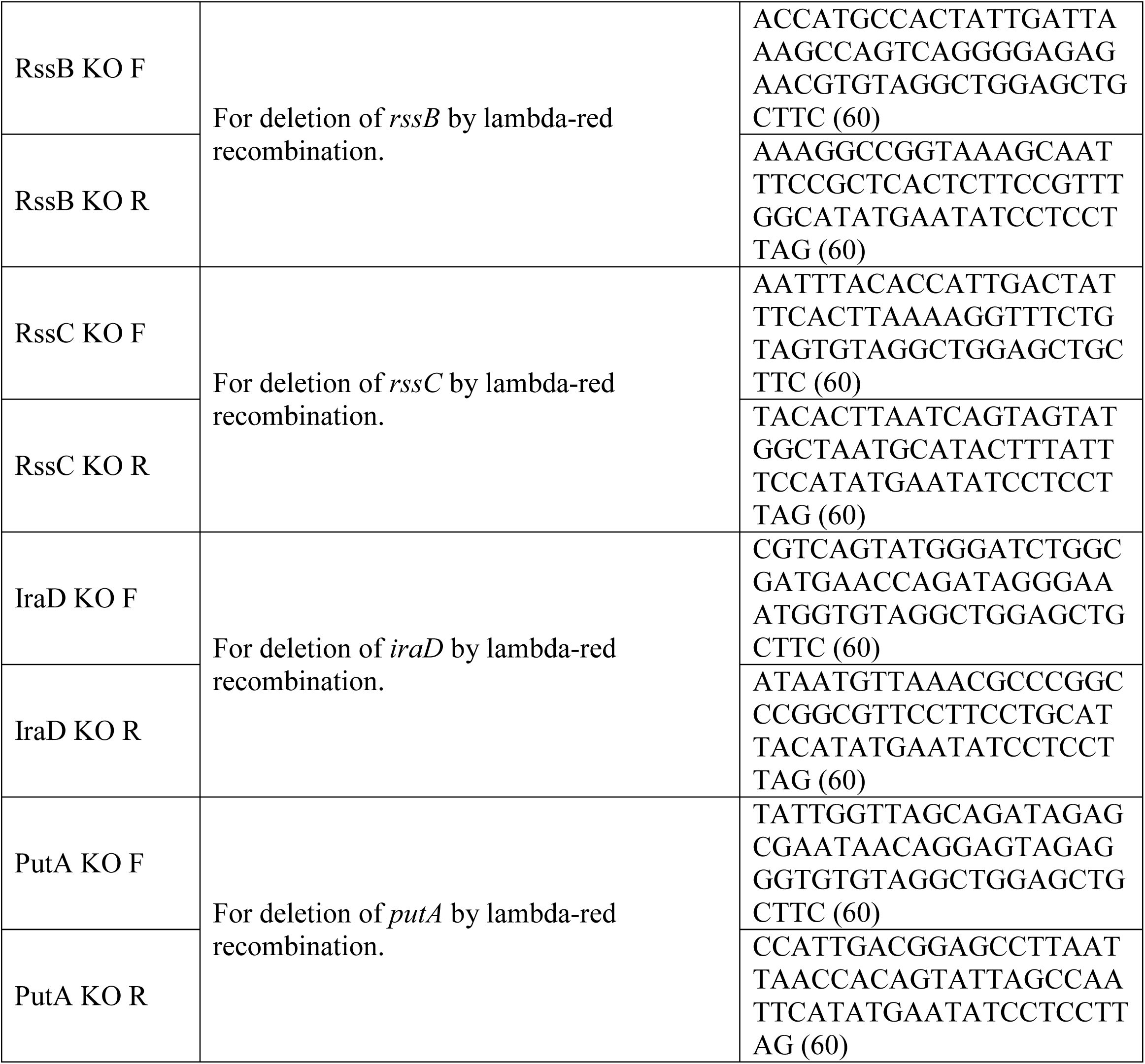
Primers

